# Spectral STED microscopy improves spectral sensitivity with polarity-sensitive probes and enables correlative measurements of membrane order and anomalous lipid diffusion

**DOI:** 10.1101/2025.02.06.636942

**Authors:** Iztok Urbančič, Falk Schneider, Silvia Galiani, Erdinc Sezgin, Christian Eggeling

## Abstract

Molecular plasma membrane organization and dynamics play an important role in cellular signalling. Advances in our understanding of the nanoscale architecture of the plasma membrane heavily rely on the development of non-invasive experimental methods, particularly of advanced fluorescence microscopy and spectroscopy techniques with high spatio-temporal resolution and sensitivity to local molecular properties. However, it remains difficult to combine several of them for a multimodal characterisation that would reduce the possibility of misinterpretations. Here, we integrated a spectral detector into a super-resolution stimulated emission depletion (STED) microscope, achieving three goals. First, we show that compared to the standard ratiometric detection using fixed bandpass filters, the spectrally resolved acquisition together with spectral fitting or phasor analysis improves the accuracy of experiments determining membrane lipid order with polarity-sensitive probes multifold. Secondly, we demonstrate that this acquisition scheme allows the use of such probes in combination with other dyes with overlapping spectra, enabling co-localisation of the membrane order maps with other cellular structures of interest, e.g. fluorescently labelled proteins. Finally, we correlate the obtained membrane lipid order with the anomalous trapped diffusion properties of a fluorescent sphingomyelin lipid analogue in the plasma membrane of living cells, as determined by STED fluorescence correlation spectroscopy, and highlight that some of the most apparent trapping sites locate at the boundaries of local ordered environments discernible by the introduced spectral STED microscopy. With additional measurements in model membranes and Monte-Carlo simulations we conclude that for sub-100 nm ordered environments uneven probe partitioning cannot by itself explain the trapping diffusion of SM in cells.

## 1 Introduction

Over the last few decades, multiple fluorescence-based microscopy techniques and molecular probes have been developed to investigate local molecular properties^1^: polarity via probes shifting their emission spectrum^2^, viscosity and tension via lifetime-changing probes^3,4^, rates of diffusion via single-molecule tracking^5^ and fluorescence correlation spectroscopy (FCS)^6^, etc. However, the spatiotemporal resolution of the most common experimental techniques is hardly sufficient to investigate the dynamic nanoscale architecture of cellular membranes, and the details of its heterogeneity remain elusive^7,8^.

Recently, several of the above-mentioned microspectroscopic methods have been coupled to super-resolution optical modalities that surpass the diffraction-limited resolution, in particular with stimulated emission depletion (STED) nanoscopy^9^. Enhancing not only the spatial but also the temporal resolution, STED has been combined with FCS (STED-FCS) enabling sub-diffraction diffusion measurements^10,11^. STED-FCS was among the first techniques to directly detect nanoscale heterogeneities in the plasma membranes of living cells, revealed via variability of the diffusion coefficient with length-scales for various membrane-anchored molecules. Most notably, fluorescent sphingomyelin (SM) lipid analogues in cellular membranes have been repeatedly shown to exhibit the so-called “trapping diffusion”^10–13^, i.e. brief slow-downs or stops in the molecular diffusion paths due to presumably transient interaction with less mobile molecules or environments. Further studies have characterised dependencies of such trapping sites on cholesterol and actin or temporal dynamics^12–14^, but their relation to differences in membrane organization such as lipid ordering remains unclear^13,15^.

Ordered lipid membrane environments are well characterized by polarity-sensitive dyes such as Laurdan or Nile-Red derivatives, which change their emission spectrum based on the polarity and thus membrane order of the immediate lipid environment^1,16^. To more efficiently detect heterogeneity in membrane lipid order, STED microscopy compatible polarity-sensitive membrane dyes have recently been presented^17,18^. This is an important step, since the enhanced spatial resolution improves the sensitivity to detect small differences in local membrane order that are otherwise averaged out by the limited spatial resolution of conventional microscopy tools. However, also other factors might contribute to the shift in emission spectrum of these dyes^19,20^. It is therefore highly desirable to correlate the reporting of several methods from the same sample^21^, but this is often unfeasible due to method-specific instrumentation requirements and experimental constraints, e.g. cross-talk between the fluorescent probes or mismatch in timescales of acquisitions.

Addressing this challenge, we here report the coupling of two complementary readouts: diffusion properties, revealed by STED-FCS, and lipid order, reported by STED-microscopy compatible emission-shifting polarity-sensitive membrane dyes. To achieve this, we integrated a spectral detector into our STED microscope, collecting the light in 12-nm bins instead of pooling all the photons over a typically broader spectral window of a bandpass filter onto a single detector. The full spectral information offers two advantages: 1) it improves the sensitivity to small changes in the local lipid order probed by polarity-sensitive dyes, and 2) it enables co-localisation of the membrane order with a fluorescently labelled structure of interest via spectral unmixing^22^, which is otherwise prohibited due to very broad emission spectrum of the Nile-Red-based polarity-sensitive dyes. Finally, we conducted these measurements in parallel with STED-FCS-based observation of transient trapping of a fluorescent SM lipid analogue. These simultaneous and correlated recordings revealed that the trapping diffusion patterns often appeared at the borders of ordered lipid environments, especially for probes with non-homogeneous partitioning between the lipid phases such as the fluorescent SM lipid analogue. Such mechanistic understanding of the readouts of advanced microscopy methods will help us further decipher the intricate nature of the membrane at the molecular scale.

## 2 Results and discussion

### 2.1 Spectrally resolved detection improves spectral sensitivity compared to the GP approach

Variations in membrane order, or lipid packing, may arise from differences in local concentrations of different lipid species^23^ – in particular of (un)saturated lipids, SM and cholesterol, and they are one of the main drivers for organising proteins in the plasma membrane^7,22,24^ and membranes of intracellular organelles^25^. To determine the physiological roles of such heterogeneities in cellular processes, it is therefore key to be able to observe and quantitatively measure variations in the local lipid order. This has traditionally been achieved with polarity-sensitive probes, such as Laurdan^26^, Di-4-ANEP dyes^27^, or Nile Red derivatives^28^. Their large shifts in emission spectrum in environments of different polarity allowed to evaluate the degree of lipid order via the normalised ratio of intensities (*I*) detected in two detection channels at two pre-defined wavelengths (λ_LO_ < λ_HI_, and corresponding *I*_LO_ and *I*_HI_) through bandpass filters. These two intensities are often combined into the normalised ratio, also termed the generalised polarisation index *GP* = (*I*_LO_ – *I*_HI_) / (*I*_LO_ + *I*_HI_)^26,29^.

Though convenient from the instrumentation standpoint, this classical two-channel ratiometric imaging does not optimally utilise the available information. The ability to distinguish two similar spectra with the GP approach depends on the selection of wavelength bands for the two-channel acquisition (i.e. λ_LO_ and λ_HI_). To demonstrate this point, we acquired the fluorescence emission spectra *I*(λ) of Di-4-ANNEPDHQ^27^ in membrane vesicles made of the lipid POPC (1-palmitoyl-2-oleoyl-glycero-3-phosphocholine) with 10%, 40% and 50% of cholesterol (Figure 1A) with a spectrofluorometer, binned the readouts in 10-nm bands, and calculated the GP values for all combinations of λ_LO_ and λ_HI_. Since the ability to distinguish any two spectra depends on detection noise, we repeated the analysis and each time added newly generated Poisson noise to the values of *I*(λ) (Figure S1A), and from these values obtained distributions of GP values for each combination of λ_LO_ and λ_HI_. We evaluated the ability to distinguish the two spectra by weighing the difference in the distributions’ mean values (*GP_i_*) with their standard deviations (*σ_GPi_*), which we call the *Contrast* = |*GP*_1_ – *GP*_2_|/(*σ_GP_*_1_ + *σ_GP_*_2_). We considered the spectra distinguishable above noise for contrast values above 1, as in the standard (z-) score (for details, see Methods and Figure S1). From Figure 1B, it is evident that the most commonly used combination of λ_LO_ and λ_HI_, typically determined by spectral peak position of the dye in pure liquid ordered and disordered (Lo/Ld) phase of model membranes (grey dot in Figure 1B), provided markedly lower sensitivity compared to the optimal pair of λ_LO_ and λ_HI_ (black dot in Figure 1B). However, for other measurements (e.g. comparing membranes with 10% and 40% cholesterol) the optimal set of λ_LO_ and λ_HI_ would be different, which cannot be known ahead or adjusted during the classical two-channel experiment.

**Figure 1:**
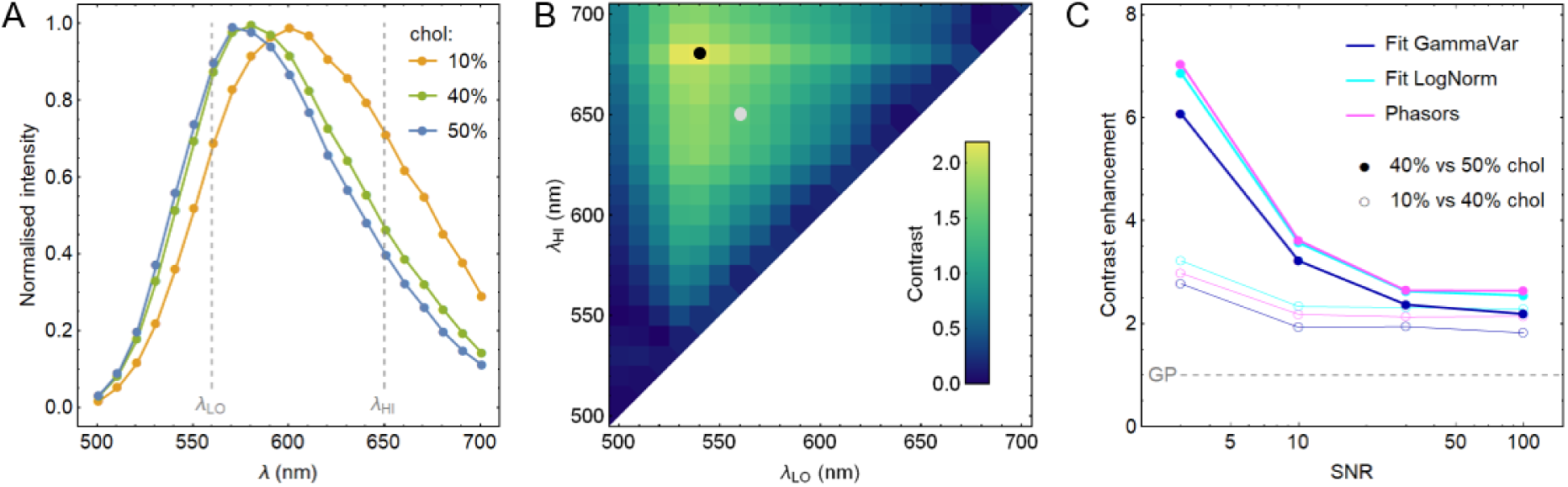
Comparison of sensitivity of two-channel and spectral detection in combination with polarity-sensitive dyes. (A) Spectra of Di-4-ANNEPDHQ in vesicles composed of POPC with variable concentration of cholesterol (chol; see colour legend), acquired with a fluorescence plate reader. (B) Simulated ability (Contrast) to distinguish the two similar spectra in panel A (40% and 50% chol) with the two-channel GP approach for different combinations of wavelengths for the two-channel acquisition (λ_LO_, λ_HI_); the dots indicate the standard combination (grey, also denoted by vertical dashed lines in panel A) and the one that gives the best contrast (black) for this pair of spectra. (C) Contrast enhancement is the relative ability to distinguish two spectra (empty and solid symbols for 10% vs 40% chol, and 40% vs 50% chol from A, respectively) by different spectral analysis methods (see colour legend), compared to the two-channel GP approach (grey dashed line), at different SNR levels. For further details, see Figure S1.

The above limitation can be overcome by collecting the signal within narrow spectral windows across the entire emission spectrum (i.e., spectral imaging/detection), e.g. through a spectral detector as in our measurements. Here, the additionally detected photons, particularly those around the peak of the spectrum that are otherwise rejected in two-channel GP measurements with limited spectral bandwidths, also carry useful information, which can be harnessed by appropriate spectral analysis. To evaluate the benefits of the full spectral information over the two channel approach above (*i*), we fitted the full spectra, generated as described before, with (*ii*) a Gamma-variate distribution^30^ and (*iii*) log-normal functions^31^, or applied (*iv*) a fit-free phasor approach^32^. Each of these methods characterises a spectrum with a set of parameters (illustrated in Figure S1B: (*ii*) GP value, (*iii*) peak position and width of the spectrum, and (*iv*) real and imaginary coordinates in a unit circle), while repeated analysis for different noisy spectra yields a distribution of these parameters. In each case, we calculated the contrast from mean values and standard deviations of the respective parameters (indicated by connecting lines in Figure S1B). Finally, we compared the contrast achieved with these three methods to the contrast obtained by the standard two-channel GP analysis. Figure 1C displaying the contrast enhancements for different SNR values shows that any of the three methods that use the full spectral information distinguished similar spectra much better than the two-channel GP approach. At high SNR ratios, the improvement was 2–3-fold, while the benefits were even larger in the most challenging experimental conditions: at low signal-to-noise ratio (SNR) and for very similar spectra (solid symbols in Figure 1C correspond to the analysis of the most similar spectra in Figure 1A for membranes with 40% and 50% cholesterol).

### 2.2 Spatial and spectral resolution of the spectral STED (specSTED) acquisition

In our previous super-resolution STED microscopy experiments with polarity-sensitive dyes and two-channel GP detection^17,18,33^, the improved spatial resolution of STED allowed us to observe finer features of heterogeneity in membrane lipid ordering. As a consequence of observing a smaller area fluorescence signal from fewer molecules is registered at once compared to confocal acquisition. Unfortunately, this also reduces the signal and SNR, and thereby lowers the accuracy of distinguishing different GP values and lipid ordering levels. Considering the results above (Figure 1C), we intended to optimise the spectral sensitivity of environment-sensitive STED microscopy with the aforementioned spectral detection methods, for which we equipped our STED microscope with a 16-channel detector with 12-nm spectral windows (Figure S2).

To highlight the capabilities of this super-resolved spectral detection mode, we first checked the spectral dependence of the spatial resolution of our STED microscope^17^. For this we labelled giant plasma membrane vesicles (GPMVs) derived from HEK 293T cells with the polarity-sensitive NR12S probe, and imaged the equatorial plane of the GPMV in confocal and STED microscopy mode. For STED microscopy, the STED laser power (140 mW at the back-aperture of the objective) was chosen to achieve an approximately 2–3-fold improvement in spatial resolution^17,18^. (Note that we evaluated the resolution with an oil immersion objective 5–10 μm deep in an aqueous samples, where spherical aberrations could have broadened the effective STED observation spot for as much as 40%^34^; further improvements in the spatial resolution can thus be anticipated closer to the coverslip, when using a water immersion objective, or by employing adaptive optics^35,36^.) The effective spatial resolution of the imaging mode was determined from the thickness of the GPMV membrane in the respective images, i.e. we determined the average values of the full-width-at-half maximum (FWHM) of the line profiles through the membrane as done before^17^. To account for residual confocal contributions, the line profiles of the STED microscopy images were fitted with either a 2-dimensional (2D) Gaussian, were the width of one of the Gaussian was fixed to the FWHM value of the confocal data, or a Lorentzian distribution. The analysis process and the final results are plotted in Figure S3. Most importantly, we determined the FWHM and the corresponding improvement in spatial resolution over the confocal data for different ranges of the fluorescence emission in windows of 12 nm around changing central wavelengths between 550 nm and 680 nm. The expected 2–3-fold improvement in resolution was achieved across the whole spectral range. More interestingly, we could observe an approximately 10-20 % higher effective resolution at the red end compared to the lower-wavelength side of the emission spectrum of NR12A. This is explained by the more effective stimulated emission cross section of the more red-emitting NR12A molecules, i.e. those experiencing a more disordered environment, as detailed before^17^.

### 2.3 To gate or not to gate

It is well known that a time-gated detection improves the resolution in STED microscopy even for pulsed laser applications^37–39^. Specifically, in time-gating the registration of photons is delayed with respect to the excitation and STED laser pulse (Figure S4A), whereby the initial contribution of non-stimulated and thus non-depleted fluorescence signal, which contributes to the aforementioned residual confocal signal in the STED images, is rejected.

For polarity-sensitive dyes time-gating may have additional impacts, since their emission spectrum is gradually red-shifting during first nanoseconds after the excitation pulse due to solvent relaxation^20,40–42^, which we could follow with time- and spectrally resolved detection (Figure S4B). Consequently, on the nanosecond time scale the difference in the spectral peak position (λ_MAX_) between the probes in Lo and Ld phase is increasing with time, implying a better contrast for delayed detection. On the other hand, much less photons are emitted at later times after the excitation pulse (Figure S4C), resulting in a reduced SNR and contrast for delayed detection. Therefore, the benefit of a nanosecond delayed (or gated) detection for distinguishing differently ordered lipid environments depends on the deployed fluorescent probe and its characteristics in the different lipid environments, specifically differences in the fluorescence lifetime and solvent relaxation times.

For NR12S in GPMVs, we find that in the confocal mode, gated detection does not improve the sensitivity to distinguish the Lo and Ld phase, while in the STED mode gating away the signal within the first 0.5–1.0 ns after the excitation pulse (as applied conventionally to suppress confocal contributions and thus to improve spatial resolution) will also slightly improve the spectral sensitivity (Figure S4F).

### 2.4 specSTED allows environment-sensitive spectral unmixing

Many cell-biological applications require the investigation of local membrane polarity relative to a structure of interest, e.g. specific proteins or protein aggregates, which are fluorescently labelled and simultaneously detected. Such co-localization measurements have well been established for the polarity-sensitive dyes Laurdan and Prodan, whose narrow emission spectrum in the blue region allowed simultaneous use of another dye, e.g. in the far-red part of the spectrum^43,44^. The broad emission spectrum of environment-sensitive probes applicable in STED microscopy, such as Di-4-ANEP and Nile Red derivatives, though, at least partly overlaps with emission of most other fluorescent proteins or probes emitting in the visible part of the spectrum, which prohibits simultaneous lipid order measurements by the standard two-channel ratiometric detection. If the spectra of all instituted fluorescence reporters is constant over space and time, one may straightforwardly distinguish a multitude of spectrally overlapping dyes with spectrally-resolved detection and spectral unmixing algorithms^45–47^, which has also been applied in STED microscopy^48^. However, such spectral unmixing is more challenging for cases where fluorescent reporters like the environment-sensitive membrane dyes are involved, whose emission spectra change with environment, i.e. potentially space and time, due to differences in membrane properties. For cases like that, we, we have recently applied multicomponent spectral fitting, which allowed us to determine the lipid membrane order at the sites of membrane proteins in live T cells using the environment-sensitive probe NR12S in combination with green fluorescent protein tags^22^.

So far, we have employed such multicomponent spectral fitting only for confocal microscopy data. Here, we demonstrate this procedure also for STED microscopy. For this, we labelled the chemically induced Lo/Ld-phase separated membranes of GPMVs^49^ filled with the green-emitting green fluorescent protein (GFP) with the polarity sensitive dye NR12S, recorded spectrally-resolved STED microscopy images of the equatorial plane (Figure 2A). For the spectrum at each spatial pixel we performed the multicomponent spectral fitting^22^, where one spectrum was fixed to that of GFP and the remaining part assigned to NR12S (Figure 2B). From this spectral decomposition, we obtained the images of extracted signal intensities for NR12S and GFP (Figure 2C and D, respectively) and the spatial maps of the spectral peak position of the NR12S emission (λ_MAX_, Figure 2E). The distinct localisation of the differently assigned signal intensities to the edge (NR12S) and in the interior (GFP) of the GPMV confirmed the successful unmixing of the signal intensities in the STED microscopy mode. The variation in spectral peak position λ_MAX_ of the NR12S emission (Figure 2E) highlighted the spectral shifts between the separated Lo and Ld phases of the membrane.

**Figure 2:**
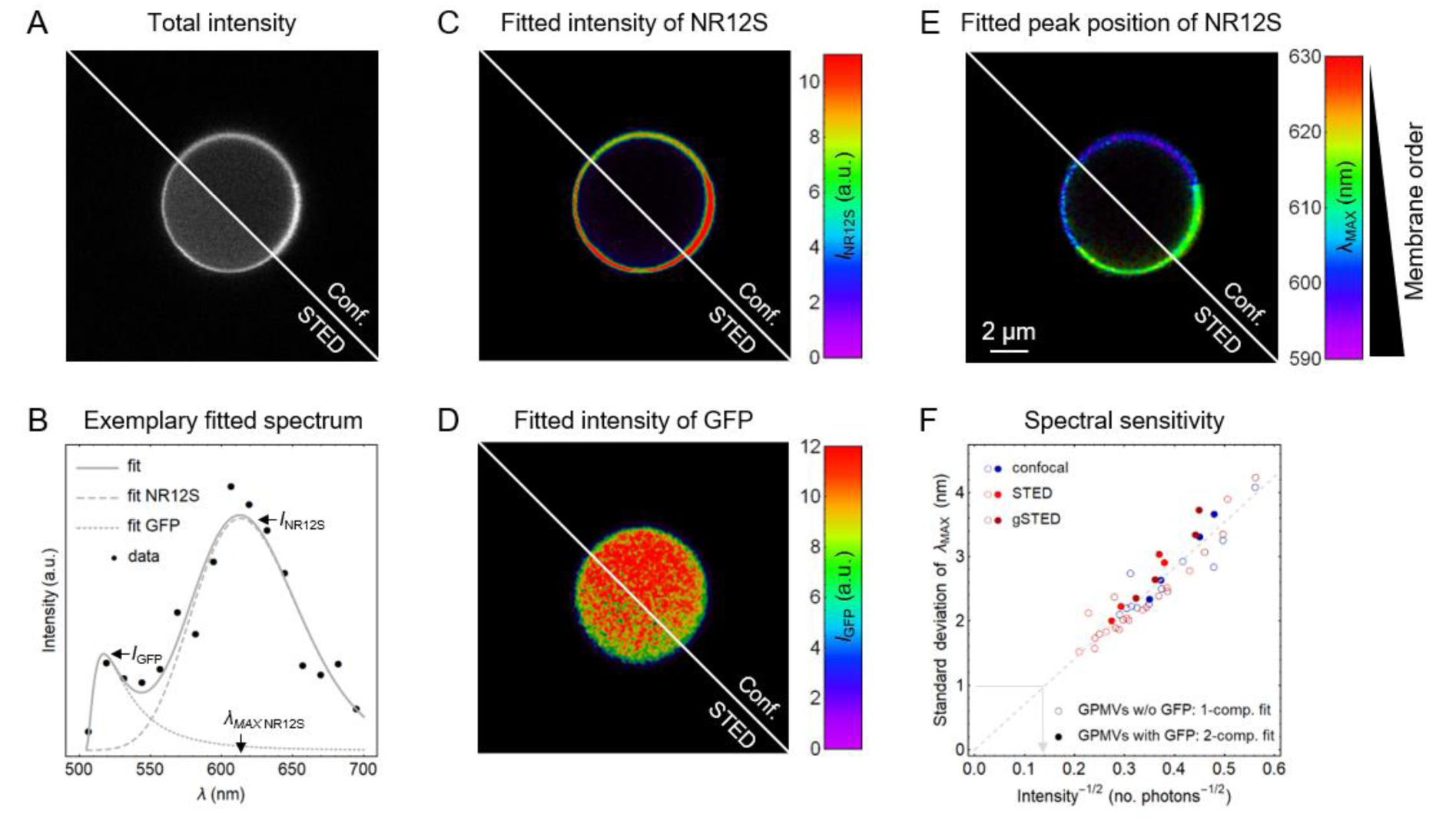
Demonstration of environment-sensitive spectral unmixing with phase-separated giant plasma membrane vesicles (GPMVs) filled with cytosolic GFP and labelled with environment-sensitive membrane probe NR12S. (A) Total fluorescence intensity image taken in the confocal and STED modes. (B) Exemplary spectrum from a single pixel (black symbols) with the two-component fit (dotted, dashed and solid grey lines represent the spectral components of the GFP and the NR12S, approximated with a log-normal function, and their sum, respectively; see Methods for details). By fitting the spectra in all pixels, we obtain the maps of fitted intensities for (C) NR12S and (D) GFP, as well as (E) the map of spectral peak position of NR12S (λ_MAX_). The clear distinction of colour along the perimeter indicates large-scale phase separation. (The shown colour scale is for confocal data; for STED it was blue-shifted for 5 nm to give a similar colorimetric impression. For absolute comparison, see for example Figure S4). (F) Standard deviation of the determined λ_MAX_ distributions for images of homogeneous (i.e. not phase-separated) GPMVs with and without GFP (solid and empty symbols, fitted with two or one spectral component, respectively), acquired in confocal, STED, or gated STED mode (blue, red, and dark red symbols, respectively), plotted against the inverse square-root of the membrane signal intensity, corresponding to SNR^−1^. Coalescence of all measurements indicates that neither the application of STED nor of the unmixing with the GFP signal affects the spectral sensitivity. The grey dashed line is a linear fit to all the data together, and the arrow indicates the estimation of the required signal intensity to obtain 1-nm spectral resolution, i.e. to distinguish two spectra separated by 1 nm.

We next checked whether the presence of the GFP signal, or the application of the STED laser for the super-resolution recordings affected the spectral sensitivity of NR12S. To this end, we imaged multiple non-phase-separated GPMVs, and determined the spectral peak position of the NR12S emission λ_MAX_ for each vesicle. Plotting the resulting standard deviation of the all values of λ_MAX_ against the inverse square-root value of the total fluorescence intensity detected for the respective GPMV, we found a linear scaling expected for Poisson noise (Figure 2F), and as described previously^31^. Notably, comparing this linear scaling for different measurement conditions, we observed no significant difference between confocal and (gated) STED microscopy measurements (blue and (dark) red symbols, respectively), neither between vesicles with GFP(solid symbols, with spectral unmixing applied) and without GFP (empty symbols, no spectral unmixing). This indicates that neither unmixing of NR12S and GFP spectra, nor the application of STED microscopy altered the sensitivity to simultaneously probed environmental cues.

Figure 2F may thus serve as a calibration for the spectral sensitivity: to resolve two spectra shifted by 1-nm, we can estimate the required fluorescence intensity (i.e., number of photons in the brightest spectral bin at that pixel) to about 50 (0.14^−2^, as indicated by the grey connecting arrow in Figure 2F).

### 2.5 Correlation of specSTED with scanning STED-FCS in live cells: trapping of SM at borders of ordered lipid sites

We finally explored the potential to combine spectral STED imaging with STED-FCS analysis of fluorescent lipid analogues. As highlighted in the introduction, our previous STED-FCS measurements revealed the so-called “trapping diffusion” for fluorescent sphingomyelin (SM) lipid analogues in the plasma membranes of living cells^10–13^, i.e. brief slow-downs or stops in the molecular diffusion paths due to presumably transient interaction with less mobile molecules or environments. Through the combination of spectral STED imaging with STED-FCS recordings we now anticipated to disclose the nature of these less mobile environments in greater detail. In STED-FCS, the apparent diffusion coefficients *D*_STED_ and *D*_conf_ are determined from FCS measurements in confocal and STED microscopy mode, i.e. for a diffraction-limited and constricted observation spots with diameters around 250 nm and 120 nm, respectively. The trapping manifests in progressively lower apparent diffusion coefficient at smaller scales (i.e. acquired at higher power of the STED laser), and is thus conveniently indicated by *D_rat_* = *D*_STED_/*D*_conf_ values below 1^10–13^. To reveal the spatial variability of the trapping, we employed scanning STED-FCS, where the observation spot is repeatedly scanned across the sample at a high frequency to yield values of the diffusion coefficient for each point along the scanned line^13^. For reliable identification of trapping sites, the recordings in confocal and STED resolution need to be performed quasi-simultaneous. For that we alternatingly scanned the same line repetitively in confocal and STED mode, as introduced in line-interleaved excitation scanning STED-FCS (LIESS-FCS)^50^. To add the information about the local lipid order, we complemented the LIESS-FCS measurement with an additional line scan, in which we excited the polarity-sensitive dye and detected its emission with the spectral detector (specSTED). We were thus repeating three different consecutive line scans: first line for confocal FCS data and *D*_conf_ determination (FCS conf), second line for STED-FCS and *D*_STED_ determination (FCS STED), and the third line for specSTED and thus determination of λ_MAX_ for NR12S (Figure 3A).

**Figure 3:**
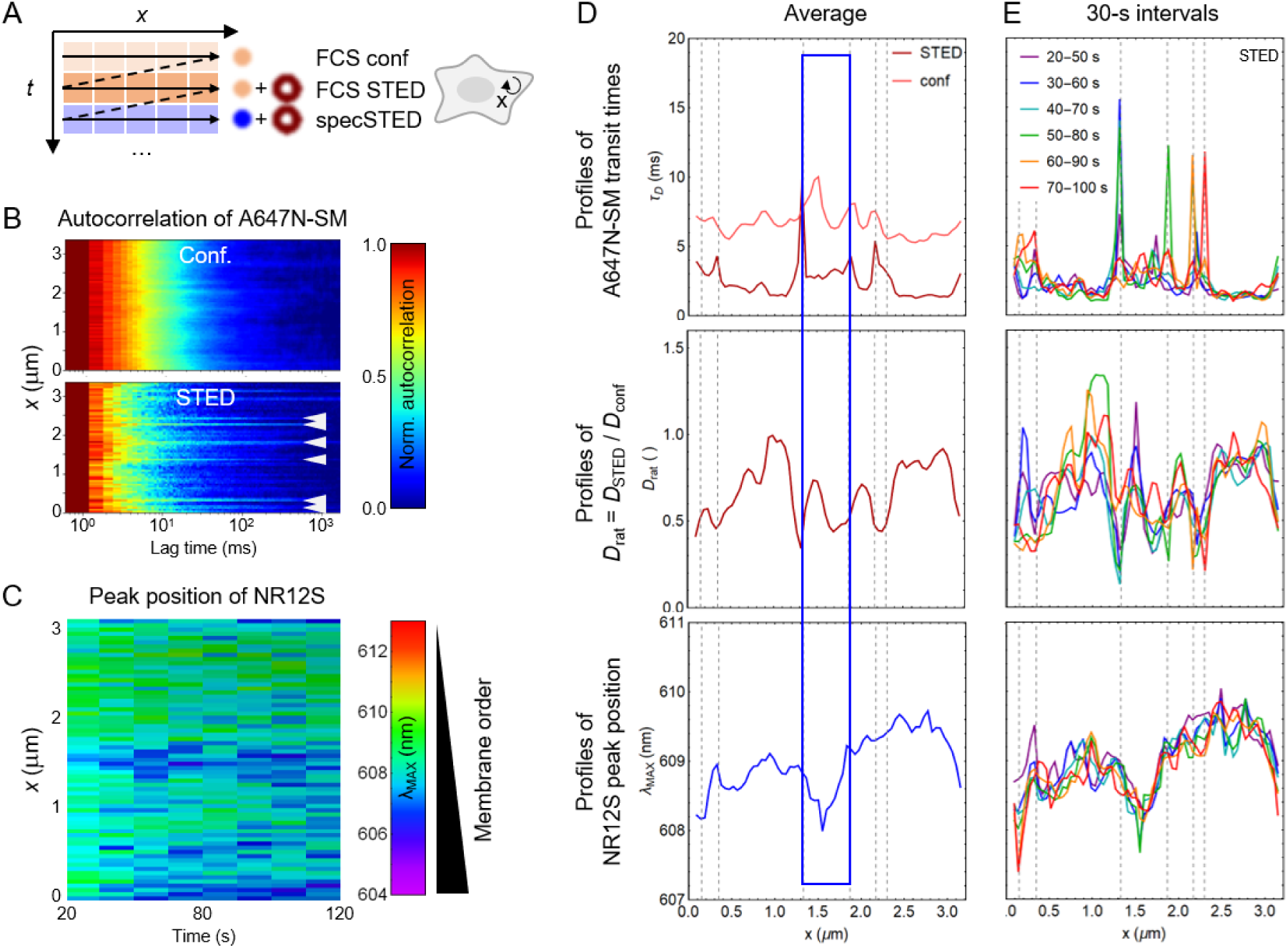
Simultaneous acquisition of scanning STED-FCS with Atto647N-SM and specSTED with NR12S in live Ptk2 cells. (A) Schematics of the measurement; the circular arrow indicates the path (*x*) for the repeated scan in the plane of the basal plasma membrane with line-interleaved STED-FCS and specSTED. The light-red and blue circles represent the activation of the 485-nm and 640-nm excitation lasers, respectively, while the dark-red ring denotes the activation of the STED laser beam. (B) STED-FCS correlation carpets, i.e. normalised auto-correlation functions plotted for every position along the scanned line, for Atto647N-SM. (B) An *xt*-profile of the spectral peak position of NR12S (λ_MAX_), binned in 10-second intervals. (D,E) Profiles of diffusion times, the ratio of STED-vs-confocal diffusion coefficients (*D*_rat_), and the NR12S λ_MAX_ (from top to bottom) calculated from the whole 100-second measurement (D) or from each 30-second segment of the data (E; colour-coded as shown). The blue rectangle indicates the region of the most pronounced area with increased lipid order (i.e. lower λ_MAX_ of NR12S), which coincides with longer diffusion times probed by STED-FCS. The vertical grey dashed lines indicate the positions of peaks in *τ*_D,STED_, identified as trapping sites (also denoted by arrows in panel B). The data indicate that the trapping sites reside at the border, rather than within the ordered lipid environments.

For our combined line-scanning STED-FCS and specSTED measurements we stained the membrane of live Ptk2 cells with the polarity-sensitive NR12S dye and the fluorescent SM lipid analogue Atto647N-SM. Figure 3B shows representative data for confocal and STED-FCS, the so-called correlation carpets (decaying FCS curves from left to right for each pixel (*x*) along the scanned line), and Figure 3C the simultaneously determined time-resolved profiles of the maximum spectral emission wavelength λ_MAX_ for NR12S along the same spatial line (*x*). Figure 3D shows the data along the scanned line averaged over a 100-second acquisition period, average transit time *τ_D_* through the observation spot as determined by confocal and STED-FCS speed (upper panel), from which we calculated the ratio *D_rat_* = *D*_STED_/*D*_conf_, which is 1 for free and below 1 for trapping diffusion (middle panel), and λ_MAX_ as a measure of local lipid membrane order (lower panel). The spatial profiles show coinciding spatial variations in lipid order and diffusion dynamics. Most notably, STED-FCS reports longer transit times *τ_D_*, i.e. slower diffusion, in the region with the most blue-shifted NR12S spectrum (denoted by the blue rectangle), which are both a result of increased lipid ordering. The values of *D_rat_* show dips to values below 1, indicating the local trapping sites often reported for SM^10–13^. However, from these data no considerable correlation between trapping diffusion and local lipid membrane ordering became obvious.

Since previous scanning STED-FCS measurements highlighted the transient nature of the trapping sites in the multiple tens of seconds time range^13,50,51^, we next cropped the measurement into 30-second intervals; the increased time resolution should avoid averaging over the appearance and disappearance of the trapping sites. Close comparison of the three simultaneously recorded data sets in Figure 3E (*τ_D_*, *D_rat_* and λ_MAX_) indicates that some of the most pronounced trapping sites (indicated by grey dotted lines) reside close to the borders of the ordered lipid membrane regions, rather than within. We have to note that we could only relate features that are stable on the scale of 30 seconds and exhibit discernible spectral variations. Although the disclosed spectral shifts were only on the order of 1 nm, we still deem them significant, as the signal intensities of about 2000 obtained for the 30-seconds segments, greatly exceeded the 1-nm sensitivity threshold as depicted in the calibration curve of Figure 2F above.

### 2.6 Monte-Carlo simulations of scanning STED-FCS: Lo partitioning produces trapping at boundaries

The observed diffusion anomaly at the borders of more ordered lipid membrane environments reminded us that fluorescent lipid analogues can show a mild or strong preference for more ordered or disordered lipid membrane environments^52,53^, which may affect also STED-FCS data^15^. To elucidate how borders between more ordered (Lo) and disordered (Ld) environments appear in scanning STED-FCS data for fluorescently labelled molecules that preferentially partition into one or the other environment, we performed Monte-Carlo simulations^54^ of the measurements with diffusing fluorescent lipid analogues that are completely or partially excluded from an ordered environment (partitioning coefficient *P_Lo_* = (concentration in Lo)/(concentration in Ld) equal to 0 and 0.2, respectively), or slightly enriched in there (*P_Lo_* = 1.2), as schematically depicted in Figure 4. We assumed a diffusion coefficient of the lipid analogue *D* = 0.5 and 0.1 μm^2^/s in Ld and Lo, respectively, which are typical values for diffusion of lipids in disordered and ordered membranes^13^ – and we did not allow molecules to exchange between the Lo environment and the surrounding area (see Supplementary Comment).

**Figure 4:**
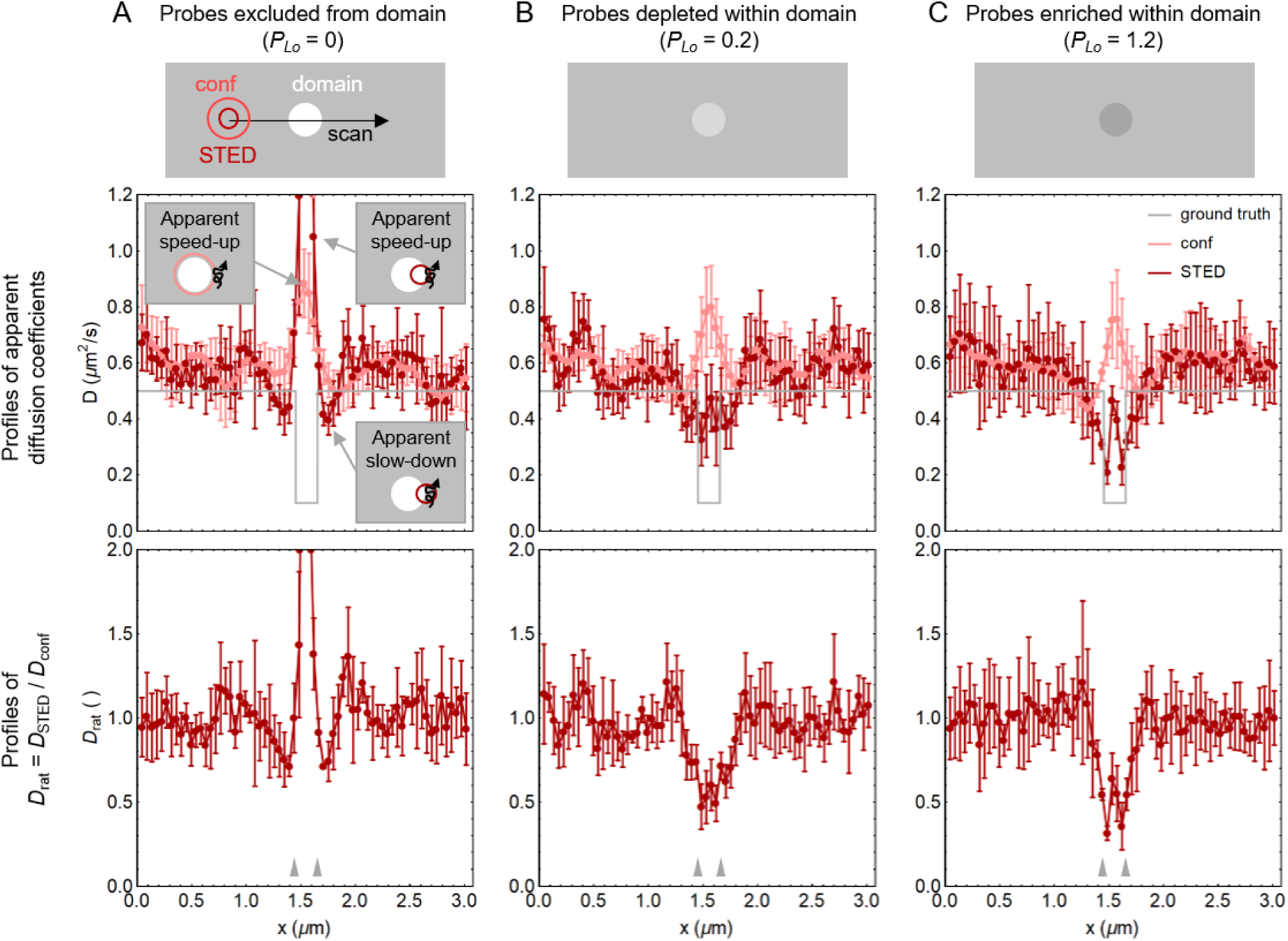
Simulations of scanning STED-FCS measurements over a lipid domain that (A) excludes the fluorescent probe, or where the probe’s concentration is (B) reduced or (C) enriched compared to the surroundings (*P_Lo_* denotes the partitioning coefficient, i.e. ratio of probe concentrations in-vs-out of the domain). Profiles of apparent diffusion coefficients (*D,* confocal and STED as light- and dark-red symbols, respectively) and their ratio (*D*_rat_) are shown in the middle and bottom rows, respectively. Datapoints and error-bars correspond to means and standard deviations of five repeated simulations. The grey lines indicate the ground-truth *D* profiles, and the arrows point to the position of the domain edges along the scanned profile. The insets within panel A sketch the origin of the variations of apparent *D* close to the domain borders. The simulated diameters of the domain, and confocal and STED PSF were 200, 240, and 110 nm, respectively.

We further chose domain-like Lo environments of size of around 200 nm in diameter (i.e., smaller than the confocal but larger than the STED microscopy observation spots) surrounded by a Ld environment, to mimic small trapping sites as highlighted before for live cells^13^. Analysing the Monte Carlo-generated diffusion tracks by line-scanning confocal and STED-FCS resulted in values of the diffusion coefficients *D*_conf_ and *D*_STED_ as well as their ratio *D_rat_* = *D*_STED_/*D*_conf_ at and outside the Lo domains (Figure 4). While the prescribed Ld diffusion coefficient of 0.5 μm^2^/s was well determined, we noted that the confocal readouts erroneously reported increased *D*_conf_ inside the domain for all three scenarios (light-red datasets in the top panels), which stems from many short transits of molecules avoiding the domain (see the scheme in the top-left inset in panel A). The amount of probe inside the domain (i.e., *P_LO_*) did not affect the confocally detected *D* profiles much, since the probes in the domain could not transit the observation spot to generate fluctuations for most of the positions along the scan.

In contrast, due to the reduced size of the observation spot the STED-FCS recordings reported different line profiles of *D*_STED_ depending on the probe partitioning. For the hypothetical case of complete exclusion of the probe from the Lo environment (*P_Lo_* = 0, Figure 4A), at positions of the observation spot close to the boundary of the domain the apparent *D*_STED_ decreased (dark-red datasets in Figure 4A), which we attributed to domain boundary preventing molecules to exit the observation spot and thereby prolonging the transit times (bottom-right inset). Closer to the domain centre, a similar-to-confocal apparent speed-up was reported. As the confocal and STED-FCS data are affected by these exclusion effects at different distances from the domain centre, their ratio *D*_rat_ oscillated around the domain boundaries (Figure 4A, bottom panel). The local dips of *D*_rat_, i.e. apparent “trapping sites” were especially pronounced at the edges of domains with enriched probe concentration (Figure 4C), where the confocal apparent speed-up effect was combined with the true detection of slowed-down diffusion in the STED-FCS mode. Of note, the observed effects were less pronounced for simulated domain sizes of 100 nm, and vanished in the noise for 50-nm domains – about half the diameter of the simulated STED beam (data not shown). Since most studies indicate that local enrichments of the ordered membrane environments in living cells are supposed to be smaller than 50 nm, the here-studied effect of the exclusion of the dye from such membrane areas is likely not the primary source of the often-reported SM trapping^10–13^.

### 2.7 STED-FCS in phase-separated model membranes: probe exclusion and inclusion appear as “trapping”

To experimentally verify how probe partitioning influences the profile of *D*_rat_ and thus trapping at local sites of more ordered Lo environments or domains, we performed scanning STED-FCS measurements across boundaries of Lo domains in phase-separated supported lipid bilayers (SLBs, Figure 5) with three different fluorescent lipid analogues: Atto647N-DPPE, Star Red (SR)-PEG-cholesterol, and SR-PEG-DSPE – all well-established STED-FCS probes^11^. From the intensity profiles across the domains (Figure 5, top row), we estimated their Lo partitioning coefficients in our lipid mixture to *P_Lo_* ∼ 0.05, ∼0.6, and ∼1.3, respectively, which qualitatively matches the three simulated scenarios above.

**Figure 5:**
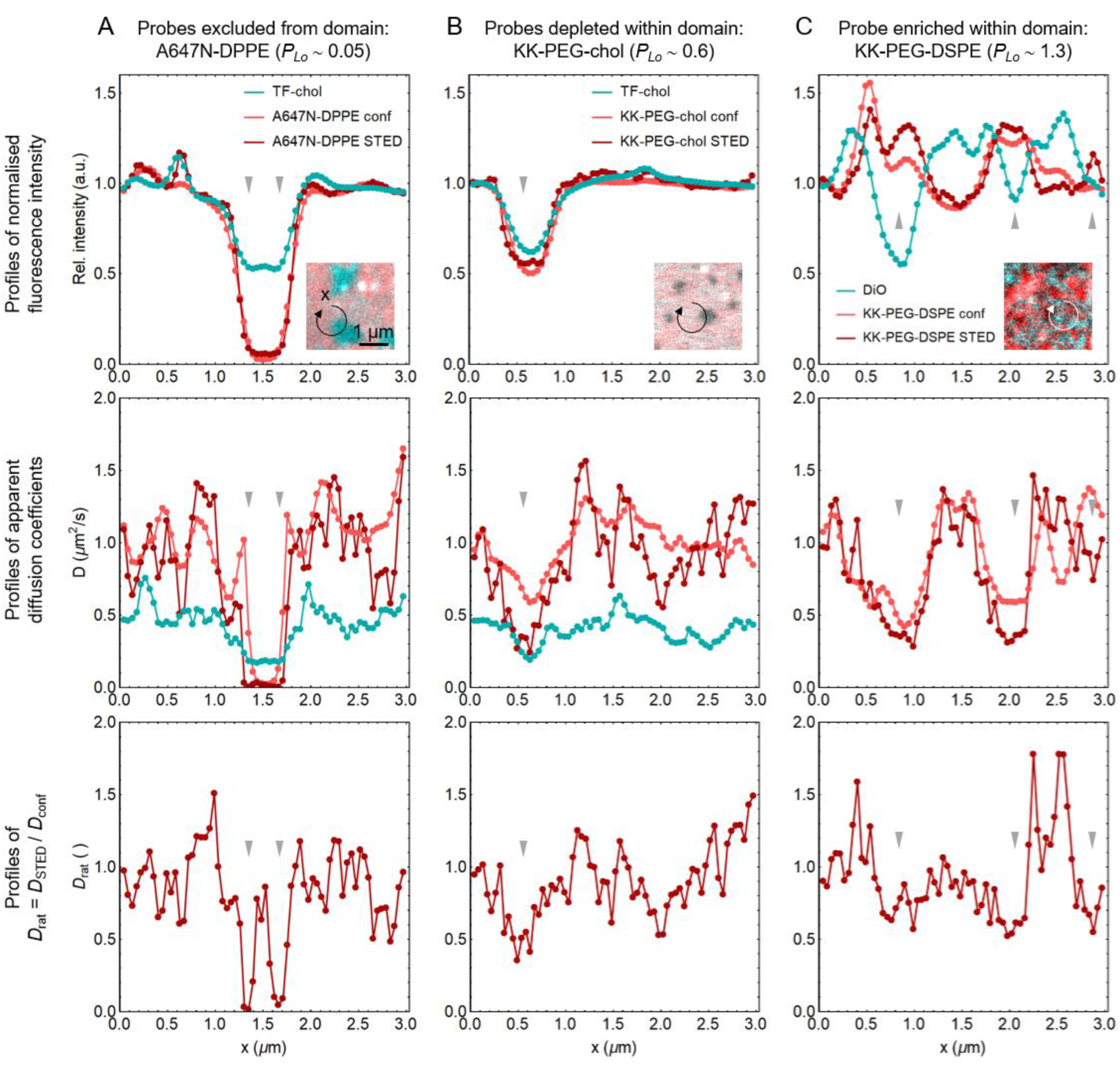
Scanning STED-FCS measurements across domain boundaries in phase-separated supported lipid bilayers (SLBs) with probes with different partitioning coefficients for the liquid-ordered lipid phase (*P_Lo_*): (A) Atto647N-DPPE is excluded from Lo, (B) SR-PEG-chol is partially depleted from Lo, whereas (C) SR-PEG-DSPE is enriched within Lo. The panels depict their normalised intensities, diffusion coefficients (*D*), and their ratio (*D*_rat_) (top, middle, and bottom row, respectively; light- and dark-red symbols for confocal and STED data, respectively). To corroborate the domain identity, another dye with known partitioning between both phases was measured simultaneously (TopFluor-cholesterol and DiO, both with slight preference for the disordered lipid phase Ld; cyan symbols). From the intensity profiles we estimated the partitioning coefficients in the Lo phase (*P_Lo_*), noted on top. The insets depict representative images of the SLBs, and circular arrows indicate exemplary trajectories of the line scans. The grey arrows indicate locations with lowest *D*_rat_. Sites for 100-s measurements were manually chosen so that the scanning trajectory crossed immobile domains.

Compared to simulations, experimental data inevitably show higher variability. Nevertheless, when the lipid analogue was almost completely excluded from the Lo phase, as was the case for Atto647N-DPPE, a pronounced “trapping site” appeared at the boundary of the Lo domain (Figure 5A; note that the data points in the very centre of Lo may not be reliable due to extremely low intensity within the domain). On the other hand, the fluorescent lipid analogues that were either only partially depleted (SR-PEG-cholesterol, Figure 5B) or even enriched (SR-PEG-DSPE, Figure 5C) within the Lo environment showed a decrease in *D*_rat_ more towards the centre of the Lo domain, as in the simulations above. Another feature common to both experimental and simulation data were regions with values of *D*_rat_ > 1 further away from domain boundaries. These resulted from probing the diffusion in the periphery of the Ld-Lo border with the two different length scales of the confocal and STED observation spots, giving another possible explanation for the sites of the so-called hop-diffusion occasionally observed in cells^50^.

## 3 Conclusions

The here-reported integration of a spectral detector into a STED microscope allowed precise co-registration of the sites of anomalous lipid diffusion characterised by STED-FCS with local variations of the membrane hydration – a proxy for local lipid order – detected by environment-sensitive fluorescent probes. For the SM analogue, known for its trapping diffusion in live cells^10,13^, the combined data showed that some of the most apparent trapping sites coincided with boundaries of ordered membrane environments.

We next aimed to check whether the observation was the direct consequence of the frustrated nature of the fluorescently tagged lipid molecule, of which the fluorescent moiety prevents the incorporation of the dye into the Lo phase, preferred by the SM part due to its interactions with lipid and protein constituents of such domains^12,55^. With STED-FCS experiments using phase-separated supported lipid bilayers and corresponding Monte-Carlo simulations we could reproduce the apparent trapping sites at the boundaries of micrometre-sized domains stable on the timescale of the acquisition (10 s– 1 min), simply due to depletion or enrichment of the probe in the ordered membrane environments. The effect stems from the sampling of molecular diffusion at different length scales close to diffusion boundaries, inherent to STED-FCS. While these observations were clear on large immobile domains of model membranes, they were much less pronounced in simulations of domains smaller than 100 nm, as well as in cells for most lipid probes except for fluorescent SM analogues – in line with previous observations^10,13^. The conclusion is consistent with the previous study with a purposefully designed Lo-partitioning probe (SR-PEG-DSPE, also used here), which would show a mildly trapping behaviour in phase-separated SLBs with domain size below 100 nm^15^, it could not recapitulate the SM-like trapping behaviour in the membranes of live cells^13^.

The stronger effect for SM-compared to DSPE-based probes could be due to the difference in the origin of their Lo preference: while DSPE probes rely on the collectivity of the weak van der Waals forces between the long saturated tails, the stronger hydrogen bonding between SM and sterols^56^ likely provide a faster-snapping interaction between the molecules during short random encounters with domain edges, before the molecules diffuse off. Overall, this work shows that uneven partitioning of the probe cannot fully explain the frequently observed sites of “trapping diffusion” for fluorescent SM analogues in live cells, for which transient stalling events really seem to be key^10^.

Though the hypothesis of distinct Lo and Ld domains^8,57^ has provided convenient conceptual framework over past decades, the reality at the molecular level likely being more complex, with domains perhaps having diffuse boundaries, similar order but different chemical compositions, etc. Local membrane curvature may further affect lipid sorting, order, and diffusion^58^. The advances of multimodal and multidimensional microscopy techniques, such as here-demonstrated specSTED with careful spectral analysis, and development of brighter and more sensitive fluorescent probes^59^, should therefore further improve characterisation of small heterogeneities in cellular membranes, and thereby elucidate the roles of lipids and membrane proteins in membrane-hosted processes such as signalling and trafficking.

## 4 Acknowledgements

The authors greatly thank Drs. Christoffer Lagerholm, Jana Koth, and Dominic Waithe from the Wolfson Imaging Centre at the MRC Weatherall Institute of Molecular Medicine, University of Oxford, for their support with imaging infrastructure, Prof. Andrey Klymchenko and Dr. Katharina Reglinski for kindly providing the probe NR12S and the GFP-SNAP plasmid, respectively, Petra Čotar for help with image analysis, and Drs. Francesco Reina and Dilip Shrestha for fruitful discussions.

The work was mainly funded by the European Commission via Marie Skłodowska-Curie individual fellowships to I.U. (grant no. 707348) and E.S. (627088), the Wolfson Foundation (No. 18272), the Medical Research Council (MRC, grant no. MC_UU_12010/unit programs G0902418 and MC_UU_12025), MRC/BBSRC/EPSRC (grant no. MR/K01577X/1), the Wellcome Trust (grant no. 104924/14/Z/14), the Deutsche Forschungsgemeinschaft (Research unit 1905 “Structure and function of the peroxisomal translocon”, Jena Excellence Cluster “Balance of the Microverse”, Collaborative Research Center 1278 “PolyTarget”), Jena Center of Soft Matter, and Oxford internal funds (John Fell Fund and EPA Cephalosporin Fund). I.U. is also supported by the Slovenian Research Agency (P1-0060 and J7-2596). E.S. is funded by Karolinska Institutet, SciLifeLab, Swedish Research Council Starting Grant (2020-02682), the Knut and Alice Wallenberg Foundation and Human Frontier Science Program (RGP0025/2022). F.S. acknowledges support from EMBO (ALTF 849-2020) and HFSP (LT000404/2021-L).

## 5 Author contributions

I.U., E.S., and C.E. conceptualised the study. I.U. and S.G. built the optical setup. I.U. carried out the experiments, analysed the data, and wrote the manuscript. F.S. performed the Monte-Carlo simulations. All authors contributed to discussions and editing of the manuscript.

## 6 Methods

### 6.1 Sample preparation: LUVs, GPMVs, PtK2 cells, phase-separated SLBs

To prepare large unilamellar vesicles (LUVs), the desired mixture of 1-palmitoyl-2-oleoyl-sn-glycero-3-phosphocholine (POPC) and cholesterol (its molar fraction was 0, 40, or 50%; both from Avanti Polar Lipids) in chloroform and methanol (1:1 vol. ratio) was dried in a glass vial, dispersed in phosphate buffer saline (PBS) by rigorous vortexing and additional bath-sonication for 5 min. The final lipid concentration was 0.2 mg/ml. The LUVs were than labelled with Di-4-ANEPPDHQ (Thermo Fisher) at a final probe-to-lipid ratio of 1:1000, and transferred into a microwell plate for measurements.

Giant plasma membrane vesicles (GPMVs) were prepared from HEK 293T cells (CRL-3216, ATCC), grown in T75 flasks (CELLSTAR) with DMEM (D5796, Sigma-Aldrich) supplemented with 10 % foetal bovine serum (FBS; Gibco), 2 mM L-glutamine, and 1 % penicillin-streptomycin (both Sigma-Aldrich), at 37 °C and 5 % CO_2_. For GPMVs filled with GFP, the cells were seeded into 35-mm cell-culturing plastic petri-dishes (Cornig), and transfected the following day with a plasmid for a GFP–SNAP-tag fusion protein (2.5 μg of DNA per dish, gift from Dr. Katharina Reglinski), together with Lipofectamine 3000 (Invitrogene), as before^60^. Next day, GPMVs were prepared following the standard protocol^61^: cells were swelled by washing in 30 % GPMV buffer (10 mM HEPES, 150 mM NaCl, 2 mM CaCl_2_; pH 7.4), followed by a 2-hour incubation at 37 °C in full GPMV buffer with dithiothreitol (DTT, 5 mM) and paraformaldehyde (PFA, 0.07 %). The harvested GPMVs were then purified by spinning down the cellular debris (2 min at 1 krpm) and concentrated by further centrifugation (15 min at 10 krpm). The membranes were labelled with NR12S at a concentration of 0.05 μg/ml for 30 min. To produce large-scale phase separation, vesicles were incubated with up to 0.5 mM deoxycholic acid (DCA). Samples were transferred into plastic-bottom ibidi μslides and imaged promptly at room temperature.

The Ptk2 cells (CCL56, ATCC) were grown in Dulbecco’s Modified Eagle’s Medium (DMEM) supplemented with 15% FBS and 1% L-glutamine (all Sigma-Aldrich). A day before the measurements, they were seeded onto 25-mm glass coverslips (#1.5 thickness by VWR). For experiments, the coverslips were mounted into metal chambers, filled with the phenol-red-free L15 medium (Sigma-Aldrich). Cell membranes we stained with both NR12S and Atto647N-SM (AttoTec) at 0.4 and 0.04 μg/ml, respectively, for 15 min at room temperature, after which the samples were washed three times with pre-warmed L15. All measurements with cells were performed at 37 °C.

Phase-separated SLBs were prepared by spin-coating the solution of the lipid mixture (1,2-dipalmitoyl-sn-glycero-3-phosphocholine (DPPC), 1,2-diphytanoyl-sn-glycero3-phosphocholine (dPhyPC), and cholesterol, in molar ratios of 1:1:0.85; all from Avanti Polar Lipids)^15^ and the STED-FCS dye (Atto647N-DPPE (AttoTec), StarRed-PEG-chol, or StarRed-PEG-DSPE (both from Abberior); approx. 1:3000 probe-to-lipid ratio) in chloroform and methanol (1:1 vol.) on 25-mm coverslips, pre-cleaned with Piranha solution^11^. The SLBs were hydrated with the SLB buffer (10 mM HEPES and 150 mM NaCl; pH 7.4). For double-checking the nature of the lipid domains, we each time added a green dye with known partitioning preference (TopFluor-cholesterol (Avanti Polar Lipids) or DiO (Sigma-Aldrich))^62,63^. The same procedure was used to prepare homogeneous SLBs composed of 1,2-dioleoyl-sn-glycero-3-phosphocholine (DOPC) and cholesterol (50:50 mol%, both from Avanti Polar Lipids), doped with fluorescent lipid analogue SR-DPPE (Abberior), used for calibration of resolution in STED-FCS.

### 6.2 Simulations of spectral sensitivity

The spectra of the Di-4-ANEPPDHQ in LUVs of different chemical composition (above) were measured with the Clariostar microplate reader within the wavelength range of 500–720 nm in 1-nm steps. To simulate spectrally resolved acquisition with a microscope, we binned the normalised spectra into 10-nm windows (Figure 1A), and added varying degrees of Poisson noise to yield a signal-to-noise ratio (SNR) of 3, 10, 30, or 100 (Figure S1A). For each condition (i.e. cholesterol concentration and SNR), we generated 4096 copies of the spectrum with independently generated noise.

We then analysed the spectra with 4 established methods: (*i*) 2-channel GP calculation^26,29^, spectral fitting with (*ii*) gamma-variate^30^ and (*iii*) log-normal functions^31^, and (*iv*) spectral phasors^32^.

i. We calculated the GP index from intensities (*I*) at two wavelengths (λ_1_ < λ_2_), with corresponding *I*_1_ and *I*_2_) according to the standard forumla^26,29^: *GP* = (*I*_1_ – *I*_2_)/(*I*_1_ + *I*_2_). Besides the “standard” combination of λ_1_ and λ_2_, we explored also all other possible combinations to find the one offering the highest sensitivity (denoted by black dots in Figure 1B and Figure S1B).
ii. For fitting of the spectrum with a gamma-variate distribution, we used the GP plugin^30^ for *Fiji*, which outputs the *GP* values calculated from the intensities of the smooth best-fit curve at the two predefined λ_1_, λ_2_.
iii. To fit the spectra with a transformed log-normal distribution, we used our optimisation routine written in *Mathematica*^31^. The parametrisation of a spectrum includes its peak position (λ_MAX_), full-width at half-maximum (*w*), and asymmetry (i.e. skewness, *a*). While *a* was fixed to a typical value for this type of probe, λ_MAX_ and *w* were subject to optimisation (Figure S1B).
iv. With the fit-free spectral phasor analysis^32^, we transformed each spectrum into a pair of real and imaginary coordinates in a unit-circle (Figure S1B):

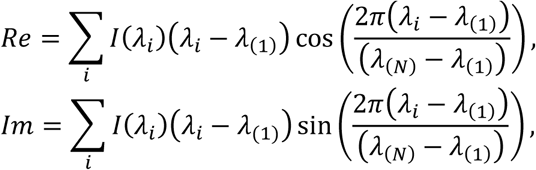

where λ_(1)_ and λ_(*N*)_ denote the wavelengths of the first and last recorded image in the λ-stack.

For each condition, we obtained the distribution of respective parameters: the GP index for *i* and *ii*, λ_MAX_ and *w* for *iii*, and the pair of real and imaginary parts of the Fourier-transformed spectrum for *iv*. We approximated each one- or two-dimensional distribution of parameters with a 1D or 2D Gaussian function, respectively (Figure S1B: smooth curves over the distributions of GP values from GammaVar fitting, or red ellipses for parameters of log-norm fitting and phasor analysis, denoting a 1-*σ* contour of the 2D fit). To evaluate the ability to distinguish two different spectra with each of the methods, i.e. the “contrast”, we calculated the distance between the corresponding parameter means Δ*p* (absolute difference for 1D parameter distributions, or Euclidean distance for 2D parameter distributions) normalised by the variability of the parameters along the line connecting the means (*σ*_Δ*i*_, denoted by thick black lines in Figure S1B): *contrast* = Δ*p*/Σ*σ*_Δ*i*_. In other words, the contrast tells the difference between the parameter means in the units of *σ*.

For the given pair of samples (i.e. 10%-vs-40% chol, or 40%-vs-50% chol, as examples of a rather large and a very small spectral difference, respectively), we calculated the contrast values at each SNR with each method (Figure S1C). We finally compared the contrast obtained with each spectral analysis method to the contrast from the direct calculation of the GP index from a standard 2-channel measurement (grey datasets in Figure S1C), and calculated the respective contrast enhancement factors (Figure 1C).

### 6.3 Experimental setup and data preparation

We used a custom STED microscope built around an Abberior Instruments RESOLFT microscope, as described before^64^ and depicted in Figure S2. In brief: the inverted body Olympus IX83 was equipped with a 100x/1.4 oil-immersion objective (Olympus) and a stage-top incubator (Okolab). The sample was excited with a 485-nm or 640-nm diode lasers (PicoQuant) pulsing at 80 MHz, steered with galvanometric scanning mirrors. For stimulated emission, the pulses from a femtosecond NIR laser (Spectra-Physics), tuned to 755 nm, were stretched by passing the light through a 20-cm-long glass rod and 120 m of single-mode polarization-maintaining optical fibre (AMS Technologies/OZ Optics). To form the ring-shaped point-spread-function (PSF) for 2D STED, we used a vortex 2π phase plate (RPC Photonics). The average laser powers, measured at the back aperture of the objective, were kept below 20 μW and 150 mW for excitation and STED, respectively.

The emitted light passed through a confocal pinhole, set to 1 Airy unit. For spectrally resolved detection, implemented within this study, we equipped the instrument with a 16-channel GaAsP PMT detector and a spectrograph (Becker&Hickl) with a diffraction grating, which split the light onto detection elements in 12-nm spectral windows. The signal from the detector, capable of multiwavelength fluorescence lifetime (specFLIM) recordings, was fed to the time-correlated singe-photon-counting unit (TCSPC; Becker&Hickl SPC150). We used this detector for the fluorescence excited by the 485-nm laser (i.e., GFP and NR12S). For classical detection of fluorescence channels in FCS, we used existing avalanche photodiodes (APD, Excelitas) and appropriate emission filters (EF): 500–570 nm (EF2 in Figure S2) for TopFluor-cholesterol and DiO, and 650–730 nm (EF3) for the far-red Atto647N- and StarRed-labelled lipids. All filters and dichroic mirrors were from Semrock or Chroma. Depending on the experiment, some of the filters and dichroic mirrors were mounted on magnetic holders and swapped as indicated in Table 1.

**Table 1:** Configuration of exchangeable elements for the three types of experiments. For their position in the optical path, see Figure S2. DM - dichroic mirror; EF - emission filter; the numbers represent the characteristic wavelengths in nm.

| Experiment | specSTED | specSTED + STED-FCS | 2-channel FCS |
| --- | --- | --- | --- |
| Element \ Detector | spectral PMT | spectral PMT + APD | 2 APDs |
| DM3 | short-pass 488 | triple-band 405/488/640 | triple-band 405/488/640 |
| DM4 | mirror | long-pass 640 | / |
| EF1 | long-pass 488 | long-pass 488 + notch 640 | / |

The microscope was controlled with the Imspector software (Abberior Instruments). The data from the spectral detector were acquired as 16-channel spectrally resolved images (i.e. λ-stack, *I*(*x*,*y*,λ)). For gating experiments, we also recorded photon arrival times (τ), binned in 0.5-ns intervals (specFLIM data: *I*(*x*,*y*,λ,τ)). To generate the gated STED data, we summed up the signal in τ channels within 0.5–8.5 ns after the excitation pulse (i.e., gate delay 0.5 ns, gate width 8 ns). For non-gated STED and confocal data, all τ channels were used. To monitor solvent relaxation (Figure S4), τ channels within 1- or 2-ns intervals were binned together.

### 6.4 Spatial resolution analysis

To determine the wavelength dependence of the STED efficiency, we acquired specFLIM data of the equatorial plane of GPMVs labelled with NR12S in both confocal and STED mode; the temporal information was used to generate the gated STED images (Figure S3A), as described above. For each GPMV image of the λ-stack, we extracted at least 50 intensity profiles along lines perpendicularly crossing the membrane, evenly spaced along the manually-drawn vesicle perimeter, using the FWHM macro in *Fiji*^17^, written by Dr. Dominic Waithe. As the signal from a single-pixel-wide profiles were extremely noisy, we averaged together 5 neighbouring profiles. Using custom *Mathematica* scripts, we approximated the confocal profiles with a Gaussian function (as exemplified Figure S3B), while the STED profiles were fitted with a Lorentz function, or with a sum of two Gaussians, where the width of one component was fixed to the value obtained from the corresponding confocal data (Figure S3C). Datapoints with unreliable fits (*R*^2^ < 0.95) were discarded. The median full-width at half-maximum (FWHM) per vesicle was used in the final display, aggregating data across 18 vesicles (Figure S3D). Finally, the STED resolution enhancement factor (Figure S3E) was calculated as the ratio of confocal and (gated) STED FWHM values.

### 6.5 Spectral fitting and environment-sensitive spectral unmixing

We used our spectral analysis software, written in Mathematica^31^. Before fitting, the background spectrum from a close-by region close to the object of interest (i.e. GPMV) was subtracted. Moving averaging of spectra across 3×3 neighbouring pixels was applied to improve the signal-to-noise ratio and thereby increase spectral sensitivity.

Each lineshape was described with a transformed log-normal spectral function^31^, parametrised by the spectral peak position (λ_MAX_), FWHM (w), and asymmetry (*a*): *S*(λ|λ_MAX_,*w*,*a*). To fit the NR12S spectra, we allowed the algorithm to optimise the λ_MAX_, while *w* and *a* were kept fixed.

For spectral unmixing of GFP and NR12S, the intensity in every pixel of the λ-stack (*I*(λ)) was fitted with a weighted sum of two spectral components (Figure 2B), as described before^22^. The parameters of the GFP spectrum, as well as the *w* and *a* of the NR12S spectrum, were predetermined from separate experiments with individual fluorophores, ensuring convergence of the unmixing algorithm by varying the remaining two parameters – the relative intensity fraction of the GFP signal (*p*) and the λ_MAX_ of NR12S:

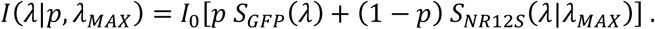

Note that the overall intensity (*I*_0_) was calculated by least-squares minimisation^31^, and is therefore not a free fitting parameter. The resulting maps of fitted parameters – intensities of GFP and NR12S (*I*_GFP_ = *I*_0_ *p*, *I*_NR12S_ = *I*_0_ (1 – *p*)), as well as λ_MAX_ of NR12S – were represented as colour-coded images (Figure 2C–E).

To further inspect the spectral sensitivity under different experimental conditions (Figure 2F), pixels in regions with membrane signal above a threshold (a third of the maximum) were selected to calculate the standard deviation of λ_MAX_ and their mean intensity.

### 6.6 Correlative specSTED and STED-FCS recordings in PtK2 cells and phase-separated SLBs

To characterise the spatial variability of diffusion properties and correlate it with variability in lipid order, we performed the measurements at manually chosen sites of the basal membrane with homogeneous intensity (flat, without vesicular structures or other bright features). For simultaneous acquisition of scanning STED-FCS data with fluorescent lipids and emission spectra of environment-sensitive membrane probe NR12S, we expanded the line-interleaved scanning STED-FCS (LIESS-FCS) approach^50^. As in scanning FCS, the observation spot was repeatedly scanned along a circular trajectory with a diameter of 1 μm, divided into 67 spatial pixels each covering approx. 47 nm of the path. For each repeat of the scan along the line, the excitation and detection configuration switched between three modalities (Figure 3A): confocal FCS (excitation 640 nm, detection with APD), STED-FCS (excitation 640 nm, STED, detection with APD), and specSTED (excitation 485 nm, STED, detection with spectral PMT). The repetition frequency of the scan was typically 2940 Hz (pixel dwell time 5 μs), or effectively 980 Hz for each modality. The total duration of the acquisition was 100 s. The resulting intensity carpets *I*(*x*,*t*) and *I*(*x*,*t*,λ) were saved as tif files for further processing.

We analysed the confocal and STED-FCS data as established previously^11^, using the freely accessible FoCuS_scan software^54^. In brief: the pixels within the first 20 s of the intensity carpets, typically contaminated by the photobleaching of immobile probe, were cropped away. When required (Figure 3E), the remaining intensity carpets were further cropped into 30-s intervals. The obtained time traces of the detected fluorescence intensity from each spatial location were then auto-correlated, yielding the so-called correlation carpets. Each autocorrelation curve was fitted with a standard model for 2D anomalous diffusion^54^, with the anomalous factor fixed to 0.9. The resulting transit times (τ_xy_) were converted into diffusion coefficients (*D* = *d*^2^/(8 ln(2) τ_xy_)). The required diameter of the PSF (*d*, measured as FWHM) of the confocal mode (240 nm) was determined with imaging of sub-diffraction-sized fluorescent beads, whereas the one for STED (110 nm) was calibrated by scanning STED-FCS experiments using the freely diffusing fluorescent lipid analogue SR-DPPE homogeneous SLBs. To characterise the diffusion modes, the ratio of the co-registered profiles of apparent *D* in STED and confocal modes (*D*_rat_ = *D*_STED_/*D*_conf_) was calculated, in which “trapping sites” appear as local values well below 1. The same procedure was used also to analyse LIESS-FCS data of fluorescently labelled lipids in phase-separated SLBs (Figure 5).

To analyse the corresponding spectral dataset, the signal was binned in 10-s intervals. The average spectrum from the first 20 s, during which the 485-nm laser was off, was considered background and thus subtracted across the whole measurement, and then cropped away. Spectral fitting was performed as described above.

### 6.7 Monte Carlo simulations of STED-FCS across a domain

The simulations of LIESS-FCS data were based on previous implementations observing freely diffusing molecules^50^. Into the standard 4 × 2 µm^2^ simulation box with reflecting boundaries, we here added a square domain of a desired size (Figure 4 shows data for a 200-nm domain). 1000 molecules of fluorescently labelled lipids were randomly seeded into the simulation box outside the domain. The number of molecules initialised inside the domain was based on a predefined partitioning coefficient *P* = *c*_in_/*c*_out_, where *c*_in_ and *c*_out_ are the number of molecules per area inside and outside the domain, respectively. We tested the following values of *P*: 0, 0.2, and 1.2, describing an empty domain, and weak or strong partitioning into the domain, respectively. Molecular tracks where simulated assuming Brownian motion and step size calculated based on the diffusion coefficients *D*_out_ = 0.5 µm^2^/s and *D*_in_ = 0.1 µm^2^/s outside and inside the domain, respectively. For simplicity, the molecules were not allowed to cross the boundaries of the domain. Together with the simulation of approx. 1000 molecular diffusion tracks, LIESS-FCS measurements were simulated on top, with settings mimicking the above experiment: a 3-µm-long scanning line with 67 pixels was placed into the centre of the simulation box crossing the domain. The fluorescence fluctuation data at every position along the scanned line were generated by calculating the overlap of the tracks with the corresponding Gaussian-shaped observation area (FWHM of 240 nm for confocal and 110 nm for STED). The pixel dwell time (and time steps) was set to 5 µs, the scanning frequency was 2000 Hz, and the total simulation time was 30 s. The intensity carpets were saved as .tiff files and analysed as the experimental data. For every condition, simulations were repeated 5 times, and the average and standard deviation per pixel reported. All simulations were performed in Python using Cython to speed up the computationally expensive checking of boundary condition at every time step, and iPyParrallel to compute multiple tracks simultaneously. The simulation code is available on Github.

## 8 Supporting information

### 8.1 Simulations of spectral sensitivity for different analysis methods

**Figure S1:**
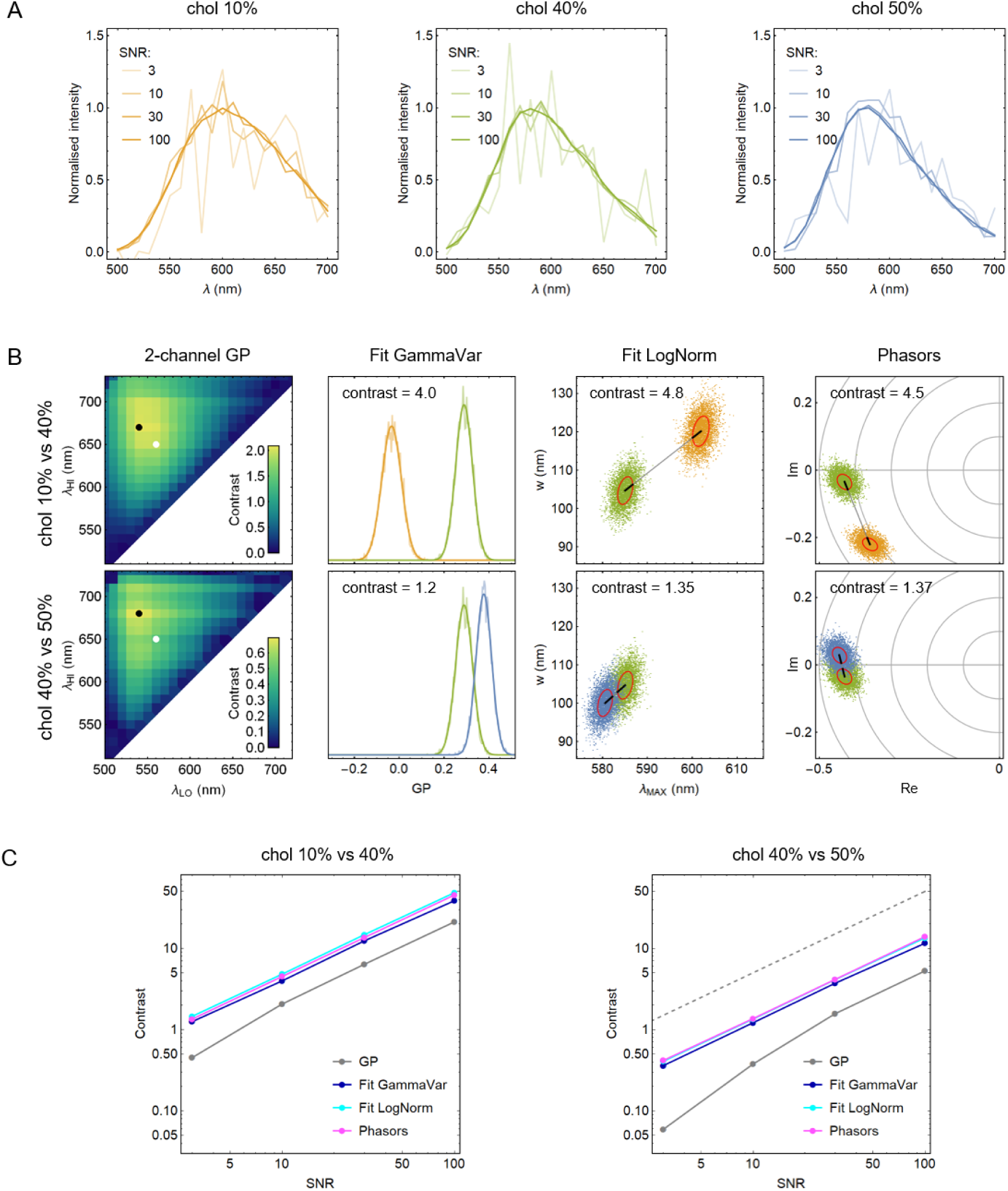
Evaluation of spectral sensitivity for different methods. (A) Examples of spectra from Figure 1A with generated Poisson noise, simulating different experimental signal-to-noise ratios (SNR). (B) For each pair of exemplary spectra (indicated in front of each row), we calculated the method-dependent contrast (spectra are resolvable if contrast > 1) after analysis of 4096 datasets with randomly generated noise (results displayed for SNR = 10); the parameter distributions were approximated by 1D Gaussians (smooth curves) or 2D Gaussian functions (allowing for xy correlations; red ellipses denote the “1 *σ*” contours); the contrast is defined as their inter-centre distance normalised by the widths along the connecting lines. (C) SNR-dependent contrast values for both pairs of spectra; the dashed line indicates the expected slope of SNR^1/2^.

### 8.2 Spectral STED setup

**Figure S2:**
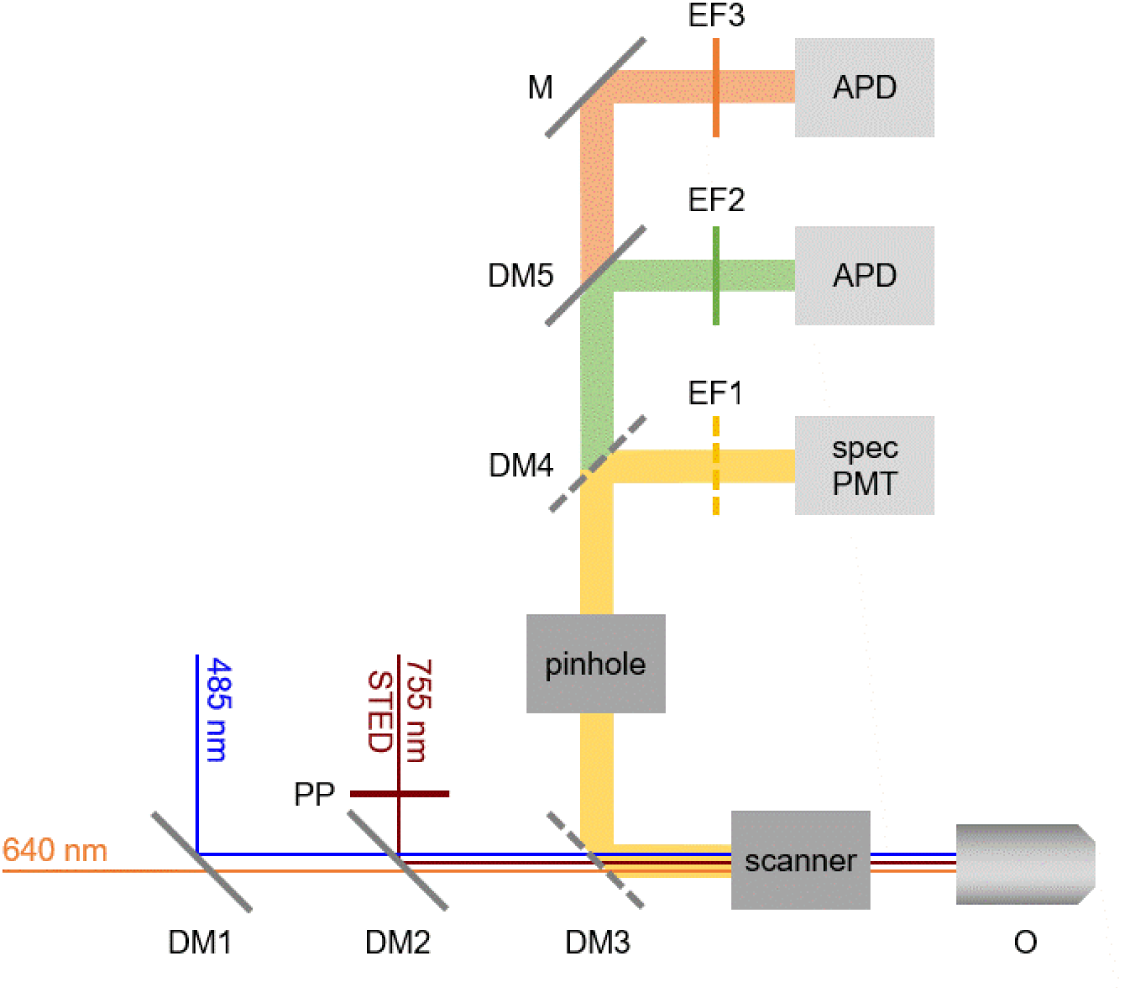
Scheme of the setup with main elements: excitation lasers (485 nm and 640 nm) and STED laser (755 nm); detection by a 16-channel array of photomultiplier tubes behind a diffraction grating (specPMT), and avalanche photodiodes (APD) for FCS; PP - phase plate; DM - dichroic mirror; M - mirror; EF - emission filter; O - objective. The dashed elements were modified for different experiments (see Methods and Table 1 for details).

### 8.3 STED resolution enhancement

**Figure S3:**
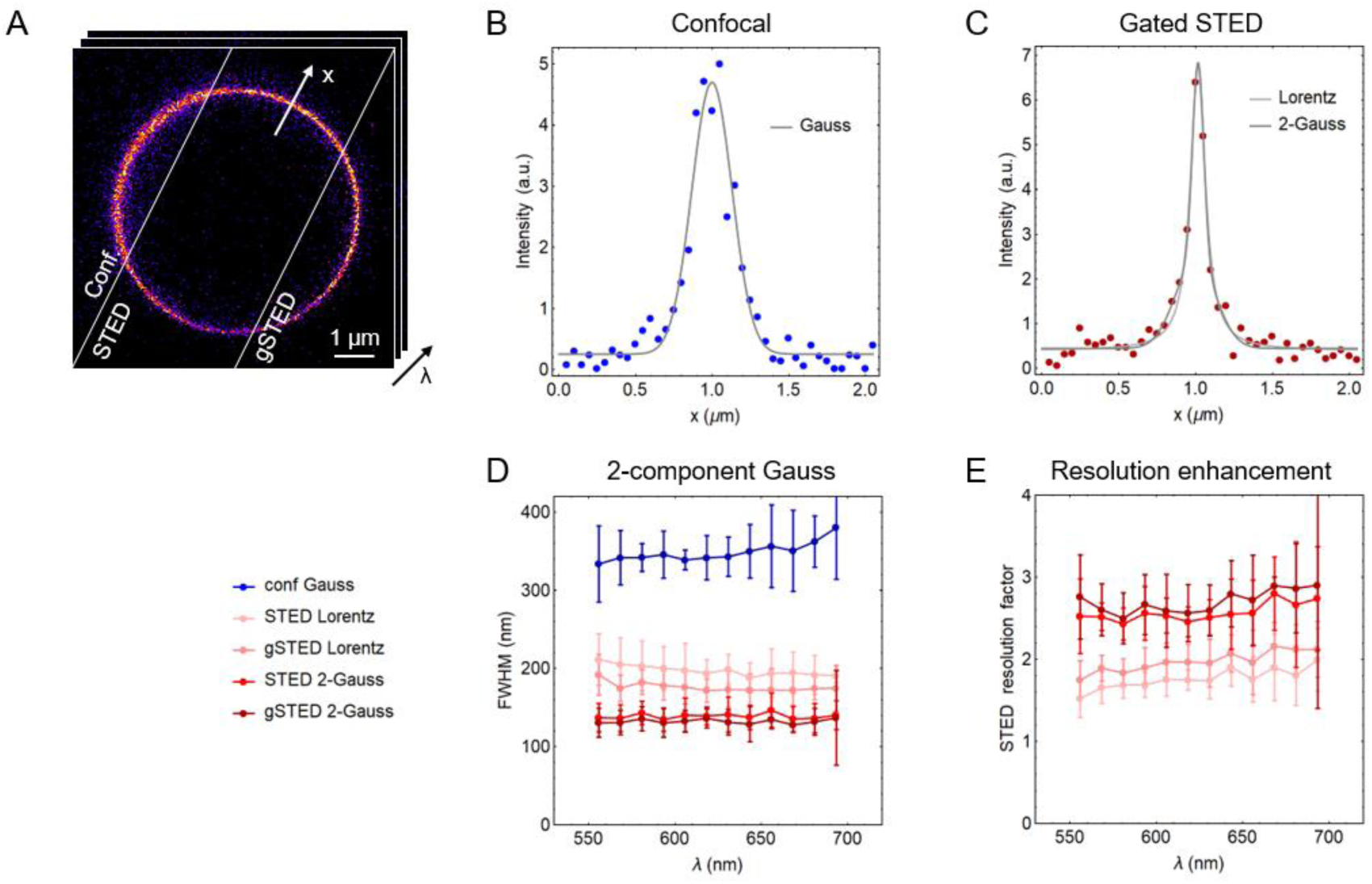
Spectral variability of the spatial resolution. (A) We evaluated the resolution via thickness (FWHM) of the intensity profiles of NR12S across the membrane in equatorial planes of giant plasma membrane vesicles (GPMVs) for spectrally resolved images acquired in confocal, STED and gated STED mode, as indicated by the arrow (*x*). The power of the STED laser was 140 mW at the back-aperture of the objective. (B,C) Exemplary intensity profiles across the membrane in confocal and gated STED (blue and dark-red symbols, respectively), together with the respective fits with a Gaussian function for confocal data, and Lorentz or two-component Gaussian function for STED data (light- and dark-grey lines, respectively). In the latter case, one component was fixed to confocal parameters to account for the contribution of the non-depleted confocal signal. (D) Spectral variability of the FWHM in all three modes, determined with different fitting approaches (see colour legend). (E) Ratio between confocal and STED resolution for data from the panel D. Symbols and error bars represent the medians and interquartile ranges for 18 vesicles.

### 8.4 Effects of gating

**Figure S4:**
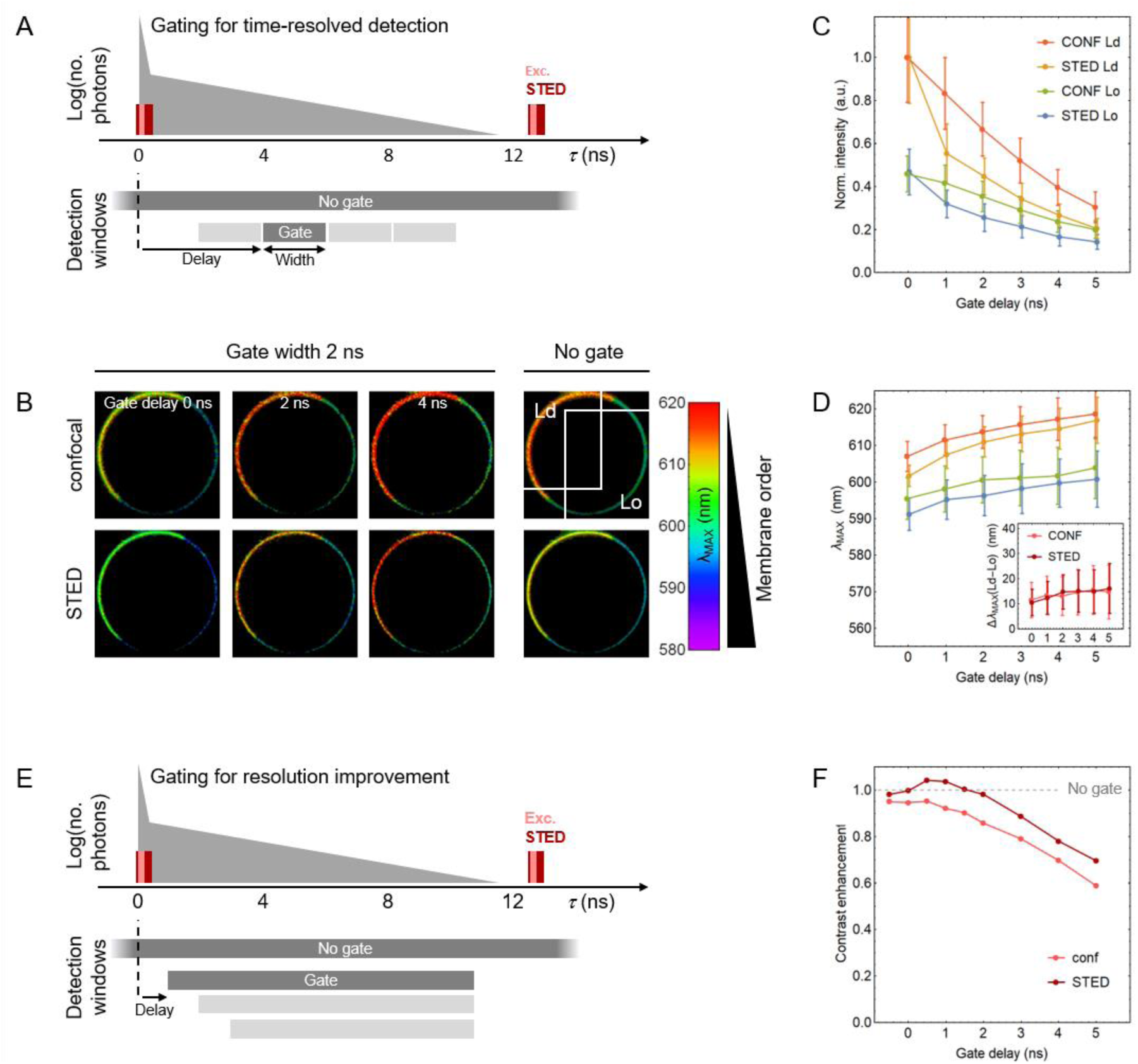
Evaluating the effect of gating in confocal and STED measurements with NR12S in model membranes. (A) Scheme of the detection windows (grey boxes) with respect to the timing of excitation and STED laser pulses (red) and emitted fluorescence photons (grey triangle) for the time-resolved detection (laser repetition rate was 80 MHz). (B) Maps of the fitted spectral peak position (λ_MAX_, colour coded as shown) of NR12S in a phase-separated GPMV in the confocal and STED mode (top and bottom rows, respectively) from data gated within 2-ns windows with different delays after the excitation pulse (left columns), or without gating (right column).From the indicated regions of identified membrane phases (Ld – liquid disordered, Lo – liquid ordered), we extracted (C) average intensities gated in 1-ns windows (normalised to Ld), showing the characteristic fluorescence decay and additional rapid initial drop due to the depletion pulse for the STED data, and (D) average λ_MAX_, the gradual increase of which stems from the solvent relaxation dynamics. Though the STED data show their emission spectrum blue-shifted for about 5 nm compared to confocal, the difference between the Lo and Ld phases (inset) is the same for both, which is consistent with previous GP analysis^17^. The error bars represent the standard deviations of the values from approx. 1000 analysed pixels across the image. (E) Scheme of the detection windows for gated detection aiming at suppression of the confocal signal. (F) Evaluation of the improvement of the spectral contrast (ability to resolve the two lipid phases, as in Figure 1C) relative to non-gated detection (i.e. value of 1, dashed grey line), with different gate delays; while gating does not improve confocal measurements, delaying detection by 0.5–1.0 ns slightly enhances the STED data.

### 8.5 Monte-Carlo simulations

#### Supplementary comment

As mentioned in the text, the probes in our simulations could not cross the domain boundaries. Though this seems like a harsh restriction, its influence on the simulated values of the transit times may not be detrimental after all. For unevenly partitioning dyes, most of the molecules hitting the border from the high-concentration side will effectively revert to keep the concentration on the other side low - hence the model approximation is reasonably valid. For an equally partitioning dye, there is equal chance of it crossing the border or bouncing off from either side. In the real case, one measures the average transit time of 3 scenaria: the fastest transits of molecules reflecting from the side of Ld, slowest transits of molecules reflecting from Lo, and mixed cases with intermediate transit times. In our simulations with no transits we only allow the first two scenaria, but as these cover both the fastest and slowest transit times of the real case, we assume that their average should approximately reflect also the realistically measured values.

## References

1. Collot, M., Pfister, S. & Klymchenko, A. S. Advanced functional fluorescent probes for cell plasma membranes. Curr. Opin. Chem. Biol. 69, 102161 (2022).

2. Klymchenko, A. S. Solvatochromic and Fluorogenic Dyes as Environment-Sensitive Probes: Design and Biological Applications. Acc. Chem. Res. 50, 366–375 (2017).

3. López-Duarte, I., Truc Vu, T., Izquierdo, M. A., Bull, J. A. & Kuimova, M. K. A molecular rotor for measuring viscosity in plasma membranes of live cells. Chem. Commun. 50, 5282–5284 (2014).

4. Colom, A. et al. A fluorescent membrane tension probe. Nat. Chem. 2018 1011 10, 1118–1125 (2018).

5. Yan, R., Chen, K. & Xu, K. Probing Nanoscale Diffusional Heterogeneities in Cellular Membranes through Multidimensional Single-Molecule and Super-Resolution Microscopy. J. Am. Chem. Soc. 142, 18866–18873 (2020).

6. Sankaran, J. & Wohland, T. Fluorescence strategies for mapping cell membrane dynamics and structures. APL Bioeng. 4, 020901 (2020).

7. Sezgin, E., Levental, I., Mayor, S. & Eggeling, C. The mystery of membrane organization: composition, regulation and roles of lipid rafts. Nat. Rev. Mol. Cell Biol. 2017 186 18, 361–374 (2017).

8. Levental, I., Levental, K. R. & Heberle, F. A. Lipid Rafts: Controversies Resolved, Mysteries Remain. Trends in Cell Biology vol. 30 341–353 Preprint at 10.1016/j.tcb.2020.01.009 (2020).

9. Klar, T. A., Jakobs, S., Dyba, M., Egner, A. & Hell, S. W. Fluorescence microscopy with diffraction resolution barrier broken by stimulated emission. Proc. Natl. Acad. Sci. U. S. A. 97, 8206–8210 (2000).

10. Eggeling, C. et al. Direct observation of the nanoscale dynamics of membrane lipids in a living cell. Nat. 2008 4577233 457, 1159–1162 (2008).

11. Sezgin, E. et al. Measuring nanoscale diffusion dynamics in cellular membranes with super-resolution STED–FCS. Nat. Protoc. 14, 1054–1083 (2019).

12. Mueller, V. et al. STED Nanoscopy Reveals Molecular Details of Cholesterol- and Cytoskeleton-Modulated Lipid Interactions in Living Cells. Biophys. J. 101, 1651 (2011).

13. Honigmann, A. et al. Scanning STED-FCS reveals spatiotemporal heterogeneity of lipid interaction in the plasma membrane of living cells. Nat. Commun. 2014 51 5, 1–12 (2014).

14. Schneider, F. et al. Diffusion of lipids and GPI-anchored proteins in actin-free plasma membrane vesicles measured by STED-FCS. Mol. Biol. Cell 28, 1507–1518 (2017).

15. Honigmann, A., Mueller, V., Hell, S. W. & Eggeling, C. STED microscopy detects and quantifies liquid phase separation in lipid membranes using a new far-red emitting fluorescent phosphoglycerolipid analogue. Faraday Discuss. 161, 77–89 (2013).

16. Parasassi, T., Krasnowska, E. K., Bagatolli, L. & Gratton, E. Laurdan and Prodan as Polarity-Sensitive Fluorescent Membrane Probes. J. Fluoresc. 8, 365–373 (1998).

17. Sezgin, E. et al. Polarity-Sensitive Probes for Superresolution Stimulated Emission Depletion Microscopy. Biophys. J. 113, (2017).

18. Carravilla, P. et al. Long-term STED imaging of membrane packing and dynamics by exchangeable polarity-sensitive dyes. Biophys. Rep. 1, 100023 (2021).

19. Gesper, A. et al. Variations in Plasma Membrane Topography Can Explain Heterogenous Diffusion Coefficients Obtained by Fluorescence Correlation Spectroscopy. Front. Cell Dev. Biol. 8, 767 (2020).

20. Amaro, M., Reina, F., Hof, M., Eggeling, C. & Sezgin, E. Laurdan and Di-4-ANEPPDHQ probe different properties of the membrane. J. Phys. Appl. Phys. 50, (2017).

21. Nicovich, P. R., Kwiatek, J. M., Ma, Y., Benda, A. & Gaus, K. FSCS Reveals the Complexity of Lipid Domain Dynamics in the Plasma Membrane of Live Cells. Biophys. J. 114, 2855–2864 (2018).

22. Urbančič, I. et al. Aggregation and mobility of membrane proteins interplay with local lipid order in the plasma membrane of T cells. FEBS Lett. 595, 2127–2146 (2021).

23. Sezgin, E. et al. Adaptive Lipid Packing and Bioactivity in Membrane Domains. PLOS ONE 10, e0123930 (2015).

24. Lorent, J. H. et al. Structural determinants and functional consequences of protein affinity for membrane rafts. Nat. Commun. 2017 81 8, 1–10 (2017).

25. Rayermann, S. P., Rayermann, G. E., Cornell, C. E., Merz, A. J. & Keller, S. L. Hallmarks of Reversible Separation of Living, Unperturbed Cell Membranes into Two Liquid Phases. Biophys. J. 113, 2425–2432 (2017).

26. Parasassi, T., De Stasio, G., Ravagnan, G., Rusch, R. M. & Gratton, E. Quantitation of lipid phases in phospholipid vesicles by the generalized polarization of Laurdan fluorescence. Biophys. J. 60, 179 (1991).

27. Jin, L., Millard, A. C., Wuskell, J. P., Clark, H. A. & Loew, L. M. Cholesterol-Enriched Lipid Domains Can Be Visualized by di-4-ANEPPDHQ with Linear and Nonlinear Optics. Biophys. J. 89, L04–L06 (2005).

28. Kucherak, O. A. et al. Switchable nile red-based probe for cholesterol and lipid order at the outer leaflet of biomembranes. J. Am. Chem. Soc. 132, 4907–4916 (2010).

29. Owen, D. M., Rentero, C., Magenau, A., Abu-Siniyeh, A. & Gaus, K. Quantitative imaging of membrane lipid order in cells and organisms. Nat. Protoc. 2012 71 7, 24–35 (2011).

30. Sezgin, E., Waithe, D., Bernardino De La Serna, J. & Eggeling, C. Spectral Imaging to Measure Heterogeneity in Membrane Lipid Packing. Chemphyschem 16, 1387 (2015).

31. Urbančič, I., Arsov, Z., Ljubetič, A., Biglino, D. & Štrancar, J. Bleaching-corrected fluorescence microspectroscopy with nanometer peak position resolution. Opt. Express 21, (2013).

32. Malacrida, L. et al. Spectral phasor analysis of LAURDAN fluorescence in live A549 lung cells to study the hydration and time evolution of intracellular lamellar body-like structures. Biochim. Biophys. Acta BBA - Biomembr. 1858, 2625–2635 (2016).

33. Barbotin, A. et al. z-STED Imaging and Spectroscopy to Investigate Nanoscale Membrane Structure and Dynamics. Biophys. J. 118, (2020).

34. Schneider, F. et al. High photon count rates improve the quality of super-resolution fluorescence fluctuation spectroscopy. J. Phys. Appl. Phys. 53, 164003 (2020).

35. Gould, T. J., Burke, D., Bewersdorf, J. & Booth, M. J. Adaptive optics enables 3D STED microscopy in aberrating specimens. Opt. Express 20, 20998 (2012).

36. Barbotin, A., Galiani, S., Urbančič, I., Eggeling, C. & Booth, M. J. Adaptive optics allows STED-FCS measurements in the cytoplasm of living cells. Opt. Express 27, (2019).

37. Moffitt, J. R. et al. Time-gating improves the spatial resolution of STED microscopy. Opt. Express Vol 19 Issue 5 Pp 4242-4254 19, 4242–4254 (2011).

38. Vicidomini, G. et al. STED Nanoscopy with Time-Gated Detection: Theoretical and Experimental Aspects. PLOS ONE 8, e54421 (2013).

39. Leutenegger, M., Eggeling, C. & Hell, S. W. Analytical description of STED microscopy performance. Opt. Express 18, 26417–26429 (2010).

40. Horng, M. L., Gardecki, J. A., Papazyan, A. & Maroncelli, M. Subpicosecond measurements of polar solvation dynamics: Coumarin 153 revisited. J. Phys. Chem. 99, 17311–17337 (1995).

41. Ma, Y., Benda, A., Kwiatek, J., Owen, D. M. & Gaus, K. Time-Resolved Laurdan Fluorescence Reveals Insights into Membrane Viscosity and Hydration Levels. Biophys. J. 115, 1498–1508 (2018).

42. Jurkiewicz, P., Sýkora, J., Olzyńska, A., Humpolíčková, J. & Hof, M. Solvent relaxation in phospholipid bilayers: Principles and recent applications. J. Fluoresc. 15, 883–894 (2005).

43. Dinic, J., Riehl, A., Adler, J. & Parmryd, I. The T cell receptor resides in ordered plasma membrane nanodomains that aggregate upon patching of the receptor. Sci. Rep. 5, 10082–10082 (2015).

44. Danylchuk, D. I., Sezgin, E., Chabert, P. & Klymchenko, A. S. Redesigning Solvatochromic Probe Laurdan for Imaging Lipid Order Selectively in Cell Plasma Membranes. Anal. Chem. 92, 14798–14805 (2020).

45. Neher, R. A. et al. Blind Source Separation Techniques for the Decomposition of Multiply Labeled Fluorescence Images. Biophys. J. 96, 3791–3800 (2009).

46. Fereidouni, F., Bader, A. N. & Gerritsen, H. C. Spectral phasor analysis allows rapid and reliable unmixing of fluorescence microscopy spectral images. Opt. Express 20, 12729 (2012).

47. Valm, A. M. et al. Applying systems-level spectral imaging and analysis to reveal the organelle interactome. Nature vol. 546 162–167 Preprint at 10.1038/nature22369 (2017).

48. Winter, F. R. et al. Multicolour nanoscopy of fixed and living cells with a single STED beam and hyperspectral detection. Sci. Rep. 7, (2017).

49. Baumgart, T. et al. Large-scale fluid/fluid phase separation of proteins and lipids in giant plasma membrane vesicles. Proc. Natl. Acad. Sci. U. S. A. 104, 3165–3170 (2007).

50. Schneider, F. et al. Nanoscale Spatiotemporal Diffusion Modes Measured by Simultaneous Confocal and Stimulated Emission Depletion Nanoscopy Imaging. Nano Lett. 18, 4233–4240 (2018).

51. Maraspini, R., Beutel, O. & Honigmann, A. Circle scanning STED fluorescence correlation spectroscopy to quantify membrane dynamics and compartmentalization. Methods 140–141, 188–197 (2018).

52. Baumgart, T., Hunt, G., Farkas, E. R., Webb, W. W. & Feigenson, G. W. Fluorescence probe partitioning between Lo/Ld phases in lipid membranes. Biochim. Biophys. Acta BBA - Biomembr. 1768, 2182–2194 (2007).

53. Sezgin, E. et al. Partitioning, diffusion, and ligand binding of raft lipid analogs in model and cellular plasma membranes. Biochim. Biophys. Acta BBA - Biomembr. 1818, 1777–1784 (2012).

54. Waithe, D. et al. Optimized processing and analysis of conventional confocal microscopy generated scanning FCS data. Methods 140–141, 62–73 (2018).

55. García-Arribas, A. B., Alonso, A. & Goñi, F. M. Cholesterol interactions with ceramide and sphingomyelin. Chem. Phys. Lipids 199, 26–34 (2016).

56. Yasuda, T., Al Sazzad, M. A., Jäntti, N. Z., Pentikäinen, O. T. & Slotte, J. P. The Influence of Hydrogen Bonding on Sphingomyelin/Colipid Interactions in Bilayer Membranes. Biophys. J. 110, 431 (2016).

57. Simons, K. & Gerl, M. Revitalizing membrane rafts: new tools and insights. Nat Rev Mol Cell Biol 11, 688–699 (2010).

58. Woodward, X., Javanainen, M., Fábián, B. & Kelly, C. V. Nanoscale membrane curvature sorts lipid phases and alters lipid diffusion. Biophys. J. (2023) doi:10.1016/J.BPJ.2023.01.001.

59. Ragaller, F. et al. Dissecting the mechanisms of environment sensitivity of smart probes for quantitative assessment of membrane properties. bioRxiv 2022.06.05.494874 (2022) doi:10.1101/2022.06.05.494874.

60. Barbotin, A., Urbančič, I., Galiani, S., Eggeling, C. & Booth, M. Background Reduction in STED-FCS Using a Bivortex Phase Mask. ACS Photonics 7, 1742–1753 (2020).

61. Sezgin, E. et al. Elucidating membrane structure and protein behavior using giant plasma membrane vesicles. Nat. Protoc. 7, 1042–1051 (2012).

62. Pinkwart, K. et al. Nanoscale dynamics of cholesterol in the cell membrane. J. Biol. Chem. 294, 12599 (2019).

63. Klymchenko, A. S. & Kreder, R. Fluorescent Probes for Lipid Rafts: From Model Membranes to Living Cells. Chem. Biol. 21, 97–113 (2014).

64. Galiani, S. et al. Super-resolution Microscopy Reveals Compartmentalization of Peroxisomal Membrane Proteins. J. Biol. Chem. 291, 16948–16962 (2016).

